# Influence of high energy diet and polygenic predisposition for obesity on postpartum health in rat dams

**DOI:** 10.1101/2021.09.24.461340

**Authors:** Andrea S. Leuthardt, Julia Bayer, Josep M. Monné Rodriguez, Christina N. Boyle

**Affiliations:** Institute of Veterinary Physiology, Vetsuisse Faculty University of Zurich (UZH), 8057 Zurich, Switzerland; Laboratory for Animal Model Pathology (LAMP), Institute of Veterinary Pathology, Vetsuisse Faculty University of Zurich, Zurich, Switzerland

## Abstract

It is estimated that 30% of pregnant women worldwide are overweight or obese, which leads to adverse health effects for both the mother and child. Women with obesity during pregnancy are at higher risk for developing both metabolic and mental disorders, such as diabetes and depression. Numerous studies have used rodent models of maternal obesity to understand its consequences on the offspring, yet characterization of changes in the dams is rare, and most rodent models rely solely on a high fat diet to induce maternal obesity, without regarding genetic propensity for obesity. Here we present the influence of both peripartum high energy diet (HE) and obesity-proneness on maternal health using selectively-bred diet-resistant (DR) and diet-induced obese (DIO) rat dams. Outbred Sprague-Dawley rats were selected and bred according to their propensity to gain weight. From F1 onward, dams consuming a HE diet displayed higher body weight gain during pregnancy, and HE diet had a strong effect on meal patterns. Sensitivity to the hormone amylin was preserved during pregnancy, regardless of diet. After several rounds of selective breeding, dams from generation F3 were assessed for their postpartum physiology and behaviors. We observed strong diet and phenotype effects on gestational weight gain, with DIO-HE dams gaining 119% more weight than DR-chow. A high-resolution analysis of maternal behaviors on postpartum day 2 (P2) did not detect main effects of diet or phenotype, but a subset of DIO dams showed decreased pup-related behaviors. During a sucrose preference test (SPT) on P14, all DR dams consumed at least 70% sucrose, while a subset of DIO dams preferred water. In generation F6/F7 dams, effects on gestational weight gain persisted, and we observed a main effect of phenotype of SPT, with DIO-chow dams showing the lowest sucrose preference. Both DIO and DR dams consuming HE diet had severe postpartum liver lipidosis and exhibited reduced leptin sensitivity in the arcuate nucleus at the time of pup-weaning. These data demonstrate that both diet and genetic obesity-proneness have consequences on maternal health.

## Introduction

Female reproductive health and metabolic status are strongly interconnected. Sufficient energy stores are permissive of ovulation, while pregnancy and lactation represent two metabolically dynamic periods of a female’s life when both the levels of and receptivity to gonadal, placental and metabolic hormones are changing rapidly. Paralleling the global rise in obesity, the prevalence of obesity in women of reproductive age continues to climb. It is estimated that up to 30% of pregnant women worldwide are obese [1; 2; 3], which increases the risk of short- and long-term adverse health outcome for both the mother and child [4]. Despite this, we still lack a basic understanding of how obesity during pregnancy and lactation can influence the metabolic and behavioral adaptions necessary to bring about new life [5].

Because maternal obesity is a significant predictor for childhood obesity, and increases the risk for metabolic syndrome and cardiovascular disease in the offspring later in life [6; 7; 8], the majority of research employing rodent models of maternal obesity has focused on the intergenerational consequences; a genetic predisposition for obesity, environmental factors, like diet, and their interaction contribute to this intergenerational risk of metabolic disorders [9]. And while it is documented that women who are overweight or obese during pregnancy are at a higher risk for developing both metabolic and mental disorders, including increased insulin resistance in early pregnancy, gestational diabetes mellitus, hypertension, pre-eclampsia and postpartum depression [4; 10; 11; 12], rodent models of maternal obesity have only seldomly been used to investigate the precise nature of the biological link between obesity-related factors and the increased risk for these maternal diseases.

A handful of studies have investigated how consumption of a high fat diet (HFD) can influence maternal behavioral and physiological parameters in mice and rats. When reported, these studies unequivocally show that consuming HFD during pregnancy leads to increased gestational weight gain [13; 14; 15]. While several demonstrated that HFD influenced metabolic and behavioral adaptations in the early postpartum period, these findings are not always consistent with one another [14; 15; 16; 17], and few have investigated the influence of diet over the full length of the postpartum period or in the post-weaning phase. What is also notable, is that each of these models of maternal obesity utilized HFD to promote increased fat mass prior to, during, or after pregnancy, but almost none considered an inherited, polygenic predisposition for obesity. Thus the question remains unanswered: does the metabolic state of obesity directly cause the elevated risk for these maternal disorders, or is rather the lifestyle or genetic factors, which contribute to obesity, that predispose these mothers to adverse health outcomes?

Decades of work in rodent dams has revealed numerous metabolic and behavioral adaptations essential for successful gestation of and caring for offspring, as well as the neural networks orchestrating them. Two exemplar adaptive systems involve the hormones leptin and oxytocin. The induction of gestational leptin resistance encourages positive energy balance during pregnancy, which is essential to support the forthcoming energy demands of late-pregnancy fetal growth and lactation [18; 19; 20]. Oxytocin-producing neurons, which are found in the supraoptic and paraventricular nuclei of the hypothalamus, undergo morphological and functional changes during pregnancy in order to promote parturition and lactation [21]. Oxytocin is important for the initiation of maternal behavior [22], and also acts as a controller of energy balance [23]. Yet, despite recognizing the interconnectedness of metabolic status and reproductive health, there are little data describing whether these, and other hormonal, adaptations are modified in the presence of maternal obesity.

In an effort to recapitulate both the polygenic and diet-sensitive nature of human obesity, we utilized two lines of rats, the lean diet-resistant (DR) and the diet-induced obese (DIO) rats. Similar to those originally described and developed by Levin and colleagues [24], DR and DIO rats were selectively-bred based on their propensity to gain weight when maintained on a palatable, high energy diet (HE). The strains of DR and DIO rats used in these studies were derived from outbred Sprague-Dawley rats and for this reason express greater epigenetic and genetic variability than inbred strains, and therefore mimic most instances of human obesity, which are polygenic in nature [25; 26; 27; 28]. Over 8 generations of breeding, we then assessed various physiological and behavioral parameters in the postpartum period, to test the following hypothesis: Both consumption of HE diet and a DIO phenotype will cause metabolic and behavioral aberration in the postpartum period. Our long-term goal is two-fold. First, generate and characterize a rat model of maternal obesity that incorporates polygenic susceptibility in an effort to better simulate most human cases of obesity, in contrast to models that rely solely on HFD. Second, to further use this model to investigate how the metabolic challenge of obesity during the already metabolically challenging periods of pregnancy and lactation can influence the maternal body and brain in ways that increase vulnerability for various metabolic and mental diseases in the postpartum period and beyond. Here we present the characterization of the development of this novel model of maternal obesity.

## Materials and Methods

### Selective breeding of DR and DIO rats

Newly established selectively-bred DR and DIO lines were initially derived from 50 male and 50 female outbred Sprague Dawley rats (RjHan:SD, Janvier Labs; Le Genest-St-Isle; France), using the protocols of Levin and colleagues [24]. At 8 weeks old, all rats were single-housed and provided ad libitum access to high energy diet (HE, Open Source Diets D12266B; 4.41 kcal/g with 32.5% kcal derived from fat, 51% kcal from carbohydrates, of which 50% is sucrose) for 3 weeks. The term “HE” will be used to specifically refer to this diet, while “HFD” will be used to refer more generally to high fat diets (typically between 45 and 60% from fat) used in previously published experiments. Body weight gain of each rat during the HE-challenge was ranked. For each sex, the 15 lowest weight-gaining rats were identified as obesity-resistant, and the 15 highest weight-gaining as obesity-prone. After at least one week maintenance on standard rodent chow (KLIBA 3436; 3.14 kcal/g, 4.5% fat/g), 12 obesity-resistant and 10 obesity-prone breeding pairs were created and designated as F0. The same HE-challenge and selection procedure was carried out for the F1 and F2 generations. Beginning with F3, the breeding pairs were semi-randomly selected (taking average-weight pups free of abnormalities at weaning) without prior HE exposure, and avoided the mating of rats who shared first or second-degree relatives. Based on nomenclature of Levin [24], rats from F3 and forward were designated as DR and DIO. The selective-breeding procedure is depicted in Figure 1A.

**Figure 1.**
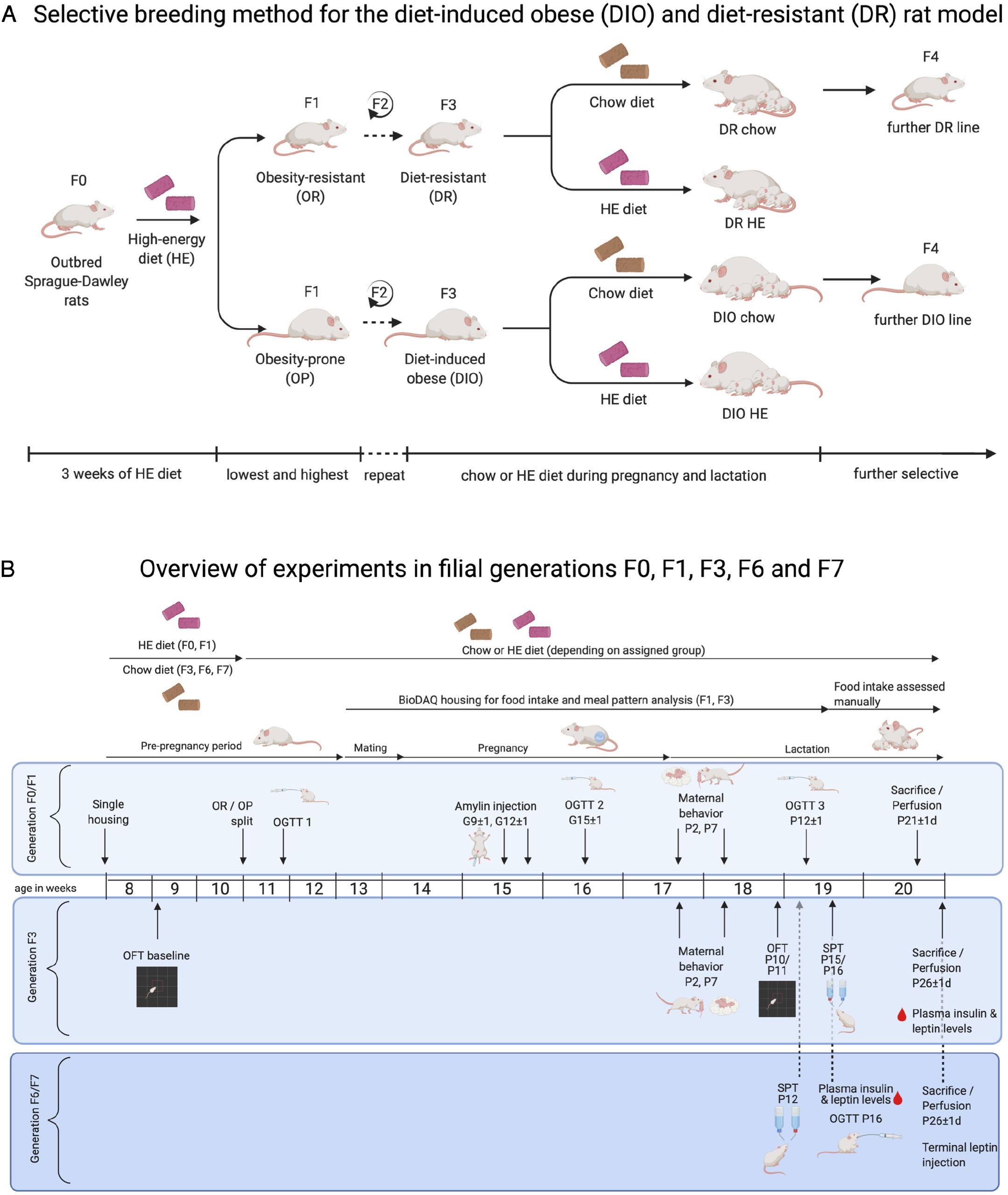
Overview of protocol for selective-breeding of DIO and DR rats and the timelines of experimental procedures in pregnant and lactating rat dams. (A) Outbred Sprague-Dawley rats (F0) were used to generate selectively-bred colonies of diet-resistant (DR) and diet-induced obese (DIO) rats, according to protocols developed by Levin and colleagues. Following a 3-week challenge on sweetened high-energy diet (HE), rats were classified as obesity-resistant (OR) or obesity-prone (OP) based on their weight gain. OR and OP male and female rats were mated, and the process was repeated in F1 and F2 offspring. Rats were designated as DR and DIO from F3 forward. For breeding and experimental purposes, F3, F6 and F7 DR and DIO dams were maintained on chow or HE during pregnancy and lactation. Offspring from chow-fed dams were used for colony maintenance. (B) Three experiments were carried out in dams from F0/F1, F3, and F6/F7 generations, respectively. Maternal body weight and food intake were collected at least weekly over the course of the studies. Experimental procedures were performed on the gestational (G) and postpartum (P) days indicated, including: oral glucose tolerance test (OGTT); amylin sensitivity test; pup retrieval and maternal behavior test; open field test (OFT); and sucrose preference test (SPT). Between P21 and P26, rat dams were perfused, and blood, brains, and livers were harvested for further analysis. Diagrams were created with a paid-subscription of BioRender.com

For some experimental groups, female dams were maintained on HE during pregnancy and lactation, however only pups derived from chow-fed dams were used to generate subsequent breeding pairs, with the exception of pups derived from F1 dams. Date of parturition was designated as postpartum day 1 (P1). In the first study dams and pups were not disturbed on P1, but in later generations dam body weight was measured on P1. On P2, maternal body weight, litter weight, size and sex ratio were recorded. Litters were then culled to 10 pups, consisting of 5 male and 5 female pups, if possible. If a dam gave birth to less than 10 pups, we cross-fostered pups from the same experimental group (matching diet and phenotype). Pups were weaned from the dam between P21 and P26.

In an effort to promote the 3R principles, the number of breeding pairs created for each generation was calculated based on the existing need for DR and DIO rats for other experiments in our laboratory. If, after assigning weanlings to colony breeding or other experiments, there was a surplus, we made every effort to re-home female rats through the Animal Welfare Office at the University of Zurich. Surplus rats that were not re-homed were killed and donated to the Zurich Zoo.

### Experimental subjects

For the present study, data were collected from rat dams from generations F0, F1, F3, F6 and F7, which were used to investigate the influences of maternal intake of HE diet and the polygenic sensitivity to this diet on maternal health parameters (described below):

#### Experiment One

Our initial comparisons of metabolic and behavioral endpoints in obesity-resistant and -prone dams from F0 (maintained on chow) and F1 (maintained on HE) yielded few differences. The first experiment therefore focused on the robust effects of HE-intake before and during pregnancy and lactation in the early generations. For the F0/F1 experiment, we therefore compared data from all chow-fed dams (n = 10) to those obtained from obesity-prone HE-fed dams (n = 14).

#### Experiment Two

The effects of the selective-breeding became more evident in F3 dams, and comparisons were made between DR-chow (n =10), DR-HE (n =5), DIO-chow (n =9), and DIO-HE (n =5) dams. This experiment focused exclusively on the postpartum period.

#### Experiment Three

Select parameters were again assessed in later generations of the DR and DIO rat dams, notably from generations F6 [DR-chow (n =8), DR-HE (n = 7), DIO-chow (n = 8), and DIO-HE (n = 6)] and F7 [DR-chow (n = 6), DR-HE (n = 6), DIO-chow (n = 6), and DIO-HE (n = 4)]. This experiment focused exclusively on the postpartum period.

An overview of the timelines for each experiment is shown in Figure 1B. All experiments were performed with the approval of the Veterinary Office of the Canton Zurich, Switzerland, and in accordance with the European Union Directive 2010/63/EU on the protection of animals used for scientific purposes.

### Measurement of body weight gain, food intake and meal patterns

For dams from the generations F0, F1 and F3, dam body weight and litter weight were recorded at least weekly, and daily between gestational day 20 (G20) and P4. Daily food intake was measured with the BioDAQ food intake monitoring system. The data was monitored with BioDAQ-E2 software (2.3.01 2011.02.10; New Brunswick; USA). For F0 and F1 dams, 24-h food intake data was analyzed to determine differences in meal patterns between pregnant (on G7 and G20 ± 1) and lactating (P4, P9, and P15 ± 1) rats, either fed with chow diet or HE. To analyze meal pattern information, the data was grouped into separate meals consisting of clustered feeding bouts. A meal was defined by a minimum inter-meal interval (IMI) of 900 sec and a minimum meal size of 0.23 g [29]. Total food intake, meal size and meal number were calculated. Following the same procedures, meal patterns of F3 dams were analyzed on G20, P4, P9, and P12. For dams from generations F6 and F7, weekly body weight was measured from date of mating until sacrifice when the pups were weaned.

### Amylin sensitivity test

In the second week of pregnancy (G11 ± 2), F0 and F1 dams were tested for amylin sensitivity. Following a 4-h fast, a single dose of amylin (20 *μ*g/kg; amylin trifluoracetate salt; Bachem, CH Catalog# 403020; Lot# 1064269) or vehicle (saline) was administered subcutaneously just before the onset of the rats’ active dark phase. Rats were tested over 2 trials in a randomized crossover manner. Food intake was measured 30 min, 1 h, 2 h, 4 h, and 24 h after injection using the BioDAQ food intake monitoring system.

### Oral glucose tolerance test (OGTT)

In F0/F1 dams, oral glucose tolerance tests (OGTTs) were performed in virgin, pregnant (G15 ± 2) and lactating (P12 ± 1) rats. Prior to the first OGTT, rats were trained to drink a 30% glucose solution from a syringe. Rats were fasted for 12 h in the light phase prior to the test. A baseline sample was obtained from tail blood, for the measurement of glucose and subsequent analysis for leptin and insulin. Rats were then given the glucose load orally (2.5 g/kg; 8.3 mL/kg consumed in 5 min), and tail blood glucose levels were measured at 15 min, 30 min, 1 h and 2 h using a glucometer (Contour XT; Ascensia Diabetes Care; CH). At each time point, a 100-*μ*l blood sample was collected for insulin determination. Blood samples were collected in Na-EDTA-coated tubes (Microvette; 100 *μ*l; Sarstedt) containing a general protease inhibitor cocktail (P2714; Sigma; 30 *μ*l of 1:10 diluted stock solution/1 ml blood). Samples were kept on ice until spun for 5 min 30 sec at 13.2 x 1000 rpm (4°C), and the plasma was immediately aliquoted and stored at −80 °C until assayed for insulin and leptin plasma levels. Total area under the curve (AUC) was calculated from 0 for glucose and insulin data using the trapezoidal rule.

OGTT was also performed in F6/F7 dams. We attempted the first OGTT in F6 dams on G15, however we observed that the pregnant DIO dams, regardless of diet, refused to orally consume the 30% glucose solution. While DR dams consumed an average of 87% of their calculated dose, DIO dams consumed 47%. Glucose (2.5 g/kg) was therefore administered via gavage for the OGTT performed on P18 ± 1 for F6 and F7 dams.

### Measurement of leptin and insulin

Leptin and insulin were measured in duplicate in plasma using the MSD® U-Plex Platform Multiplex Assay (U-PLEX Metabolic 2-Plex Combo 1 (Rat), Catalog# K15312K-2, Lot# 348956); plates were analyzed using the MESO QuickPlex SQ 120 imager. Hormones were measured in the OGTT baseline samples for F0, F1, F6, and F7 dams (after a 12-h fast), and in terminal samples for F3 (after a 2-h fast).

### Behavioral parameters

#### Maternal Behavior Tests

To assess maternal motivation, a pup retrieval test followed by 60-min of home cage observation was performed in the F0/F1 and F3 dams on P2 and P7. Maternal behavior tests were conducted in the first four hours after dark onset. Before each test, location of the pups and nest was recorded, and then pups were placed in holding cage under warming light for 10 min. The test was initiated when 5 pups were placed in each of the two corners of the home cage opposite of the nest.

For tests with F0/F1 dams, maternal behaviors were manually recorded in real-time with a stop-watch. The latency to retrieve the first pup and all pups, and the latency to assume a nursing position was recorded. Subsequently, the dam’s behavior was recorded every minute for 60 min. Each behavior was assigned to one or maximum two of the following categories: retrieving, nursing (further defined as hover, high crouch, or low crouch; See [30] for reference), pup-grooming, self-grooming, eating, drinking, exploring, or quiet (out of the nest). Using this manual scoring, the resolution of these data was at the level of minutes.

The primary aim for the F3 generation was to perform a higher resolution analysis of maternal behaviors than was obtained during F0/F1. The general timeline matched the F0/F1 maternal behavior test, but to achieve a finer detail, the 60-min test was recorded using a RaspPi Camera. Up to 6 dams were tested in a single session, and the behaviors of the dams were blindly scored using the free and open-source software Behavioral Observation Research Interactive Software (BORIS) [31]. The following behaviors were analyzed: latency to retrieve first pup, latency to retrieve all pups, nursing duration, nursing positions (hover, low crouch and high crouch), nest building, pup grooming, self-grooming, eating, drinking, exploring and quiet (out of the nest). These behaviors were sub-categorized as pup-directed or self-directed behaviors. Pup retrieval, pup grooming, nursing and nursing position accounted for the pup-directed behavior, whereas self-directed behavior included self-grooming, eating, drinking, exploring and quiet. Additional details of the analysis can be found in the Supplemental Methods. By scoring the videos using BORIS, duration of each behavior during the test could be evaluated at the level of seconds.

#### Open Field Test

In F3 dams, open field tests (OFT) were conducted in each female rat prior to mating or access to HE diet (baseline, at approximately 9 weeks of age), and again on postpartum day 10 or 11. Tests were performed during the light phase, between 8 and 11 am, in a testing room separate from the animal housing room. The OFTs were conducted in an open metal frame (80×79×40 cm). The floor of the frame was covered with a transparent plastic floor to facilitate cleaning, under which black plastic was placed to increase contrast of white rat in the frame. Dams were always placed into the same corner of the square and immediately left alone in the room. The behavior and movement of the animal was recorded for 15 minutes with a video camera (RaspPi Camera). After the test the rat was brought back to her home cage, the floor and walls of the Open Field were cleaned with Virkon™ S and water, and dried.

Videos were analyzed using ezTrack [32], a free and open source program using jupyter notebook and interactive Python scripts (Python 3.7.). All required programs used for analysis were downloaded from the Github Denise Cai Lab ezTrack website (https://github.com/DeniseCaiLab/ezTrack). Data were summarized using binned summary reports. Bins were defined as frames: 0-27000; 0-9000; 9001-18000; 18001-27000, which represent 0-15 minutes of the video, 0-5 minutes, 5-10 minutes and 10-15 minutes. For each bin the following parameters were analyzed (all data in centimeters unless defined otherwise): Total distance, total distance per bin, total periphery zone distance, total center zone distance, center zone distance per bin, percentage of total time spent in center, total number of center entries, total number of center zone entries per bin.

#### Sucrose Preference Test

Beginning at postpartum day 12, 4 consecutive days of sucrose preference testing (SPT) were conducted with rat dams maintained in their home cages. The standard water bottle (500 ml) was replaced with two smaller water bottles (250 ml each); one containing 1% sucrose solution (Sucrose ACS reagent, Sigma-Aldrich) and the other containing water. Each bottle was weighed and refilled every day. The left-right placement of the bottles was changed each day to distinguish a side preference from the actual sucrose preference. For each day of testing, any preference for or avoidance of the sucrose solution was calculated as percentage of the total volume consumed: [100 x volume of sucrose solution consumed/total volume consumed]. The intake of sucrose and water on the last two days of testing were averaged. Data from rats demonstrating a consistent side-preference (drinking 75% or more from the same side) were not included in the analysis.

### Immunohistological analysis of postpartum brain tissue

#### Perfusion and tissue preparation

Rat dams were sacrificed between postpartum days 21 and 26. On the morning of perfusion, pups were separated from dams at least one hour prior to any treatment, at which time food was removed from the cage. To test for central leptin sensitivity, dams from F3 and F7 were treated with leptin prior to perfusion. Within the experimental groups, dams were randomly assigned to receive PBS vehicle or leptin (i.p., Murine leptin; G2817 Lot: 071776; PeproTech; UK; Diluted in 0.01 M PBS, pH 8.0). Dams from F3 were treated with 2 mg/kg, but since this failed to induce pSTAT3 in the hypothalamus, F7 dams were subsequently treated with 5 mg/kg. Forty-five minutes after leptin or vehicle injection, rats were injected with an i.p. injection of 300-450 mg/kg Pentobarbital (Esconarkon ad us. Vet.; 300 mg/ml; Streuli Pharma AG; Uznach CH). When deeply anesthetized, a blood sample was taken from the heart, transferred to a Na-EDTA coated tube, and kept on ice before centrifuged for 10 minutes at 13’200 rpm. Plasma was aliquoted and directly frozen on dry ice. Following blood collection, samples were collected from the right median lobe of the liver and visceral adipose tissue; one portion of each were snap-frozen in liquid nitrogen, and one portion stored in 4% paraformaldehyde (PFA) for histological analysis. The rat was then flushed by hydrostatic forces with 0.1 M ice cold phosphate buffer (PB) for 1.5 minutes followed by a 2.5-minute fixation with ice-cold 2% PFA. The brain was extracted and put into 2% PFA on ice. Brains were transferred to 20% sucrose solution in 0.1 M PB for 24 h in a 4°C cool room. Brains were blocked into fore- and hindbrain, frozen in hexane on dry ice for 4 minutes, and stored at −80°C until sectioning.

#### Histological analysis of postpartum liver

Liver tissue samples collected from F0/F1, F3, and F7 dams were sent to the Institute of Veterinary Pathology at the University of Zurich for further evaluation. Liver tissue was embedded into a paraffin block, sliced in 3-5 *μ*m thick sections and mounted onto glass slides. Following hematoxylin and eosin-staining (H&E), slides were scanned using a slide scanner (NanoZoomer-XR C12000; Hamamatsu, Hamamatsu City, Japan). Liver sections were semi-quantitatively scored using the non-alcoholic fatty liver disease (NAFLD) Clinical Research Network Scoring System [33], taking into account that NAFLD in rodents is not associated with the development of megamitochondria and a few other features described in human NAFLD [34]. Under this lipidosis scoring system, a score of 0 represents not present, 1 represents very mild, 2 represents mild, 3 represents moderate, and 4 represents severe lipidosis.

#### Immunohistochemistry in brain

Frozen forebrains of DIO/DR generation F7 were sectioned in four series of 25 *μ*m thickness on a cryostat (Leica CM3050 S; Biosystems; DE). Sections were mounted directly onto Superfrost® glass slides (Superfrost® Plus; Thermo Scientific; DE), beginning at +0.2 mm from bregma through the hypothalamus until −3.25 mm from bregma, according to the Swanson Brain Maps Atlas [35]. Between −0.83 and −1.08 from bregma a total of 4 sections (200 *μ*m in total) were discarded. All other sections were collected. Slides were stored at −20°C in cryoprotectant solution (20 % glycerol; 30 % ethylene glycol; 50% phosphate buffer (0.1M PB)) until staining.

#### Oxytocin and pSTAT3 Immunostaining

Two series of the brain sections were separately analyzed for oxytocin and pSTAT3 expression using immunofluorescence protocol optimized for the visualization of pSTAT3 (see [36] for details). In brief, slides were blocked in 4% NDS, 0.4% Triton and 1% BSA in KPBS for 20 minutes and then incubated in rabbit-α-pSTAT3 (1:500; 9145 from Cell Signaling Technology) or mouse-α-Oxytocin (1:1000; Cat # MAB5296; Lot # 3061232; Chemicon; Temecula; CA) in 1% NDS, 0.4% Triton and 1% BSA in KPBS for two overnights at 4°C. After primary incubation slides were rinsed in KPBS, followed by a secondary incubation step in donkey-α-rabbit–Cy3 or donkey-α-mouse–Cy3 (both 1:100; Jackson ImmunoResearch, Laboratories) in 1% NDS and 0.3% Triton in KPBS for two hours at room temperature protected from light. Slides were rinsed again in KPBS, counterstained in DAPI for 4 min and washed in KPBS. After 30 min air drying slides were cover-slipped with Vectashield® (Vectashield® Hardset™; Antifade Mounting Medium for Fluorescence; Ref # H-1400; Lot # ZH0114). Slides were stored at 4°C in the dark until further analysis.

#### Fluorescence microscopy and image analysis

Fluorescence images were taken at 10x magnification for oxytocin neurons and at 20x magnification for pSTAT3 neurons with a Zeiss Axio Camera HRm (AxioCam HRm, Carl Zeiss Microimaging GmbH, 37081 Göttignen, Germany). Sections were excited at 550 nm to visualize pSTAT3- or oxytocin-positive neurons depending on the slide series.

Images were analyzed using Image J software (Version 2.0.0-rc-69/1.52p; open source image processing software; [37]) together with short macro programs customized for quantification of pSTAT3- or oxytocin-positive neurons. For analysis, 8-bit grayscale pictures were used. Threshold for pSTAT3 neuron count was set at 38-255 and the size of counted cells was defined as 30 square pixels up to infinity. For the evaluation of oxytocin cells, first the function “Gaussian Blur…” with a sigma value = 2 was applied. Threshold was then set at 40-255, size was set at 40 square pixels up to infinity and the application “Watershed” was used to divide overlapping cell bodies. For oxytocin, the macro was run on brain sections containing paraventricular nucleus of the hypothalamus (PVH, corresponding to Levels 24 through 27 in [35]). For pSTAT3, brain sections containing arcuate nucleus of the hypothalamus (ARH, Levels 28 through 30 in [35]) were analyzed.

#### Statistical analysis

For all statistical analyses Microsoft® Excel (version 16.48) and Prism 9 (version 9.1.0) were used. Depending on the dataset, mixed-effect model analysis, two-way-ANOVA or unpaired parametric two-tailed Student’s t-tests were used to evaluate the influence of diet (chow vs HE), phenotype (DR vs DIO), and time on each parameter measured. To account for missing values in a repeated measures ANOVA, a mixed-effect model analysis was used. For multiple comparisons (post-hoc tests), a Tukey’s or Sidak’s multiple comparison was used. To analyze ordinal data (semi-quantitative NAFLD scores), nonparametric Kruskal-Wallis tests were used, followed by Dunn’s planned multiple comparison test. P-values ≤ 0.05 were defined as statistically significant. Data are represented as mean ± SD.

## Results

### Experiment One: Influence of HE on obesity-prone and -resistant rat dams (F0 and F1) during pregnancy and lactation

#### Maternal body weight gain and food intake

Compared to dams eating chow, maternal intake of HE resulted in differences in body weight (Fig. 2A) and body weight gain (Fig. 2B), which were especially pronounced during gestation and the first week of lactation. Main effects of diet (F_1,22_ = 27.25, P < 0.001) and time (F_2.60,53.19_ = 294.0, P < 0.001) on body weight were found, with significantly higher body weight in HE-fed rats at all time points excluding P14 and P21 (Fig. 2A). Furthermore, a significant interaction was detected (F_15,307_ = 12.57, P < 0.001). Main effects of diet (F_1,21_ = 6.592, P = 0.018) and time (F_2.61,53.76_ = 218.8, P < 0.001) on body weight gain were found, with significantly increased body weight gain in HE-fed rats compared to chow-fed rats during early- to mid-pregnancy, and on P2, P14 and P21 (Fig 2B). Comparison of body weight gain during pregnancy (the difference between body weight on G1 and on P2 to account for litter and placenta weight; See inset in Fig. 2B) revealed that dams fed HE gained significantly more weight than chow-fed dams during pregnancy (t_21.99_ = 2.925, P = 0.008). HE also influenced the caloric intake during pregnancy and postpartum: Main effects of diet (F_1,21_ = 9.32, P = 0.006) and time (F_7.86,172.6_ = 130.2, P < 0.001) were observed, with individual differences observed in early and midpregnancy (Fig. 2C).

**Figure 2.**
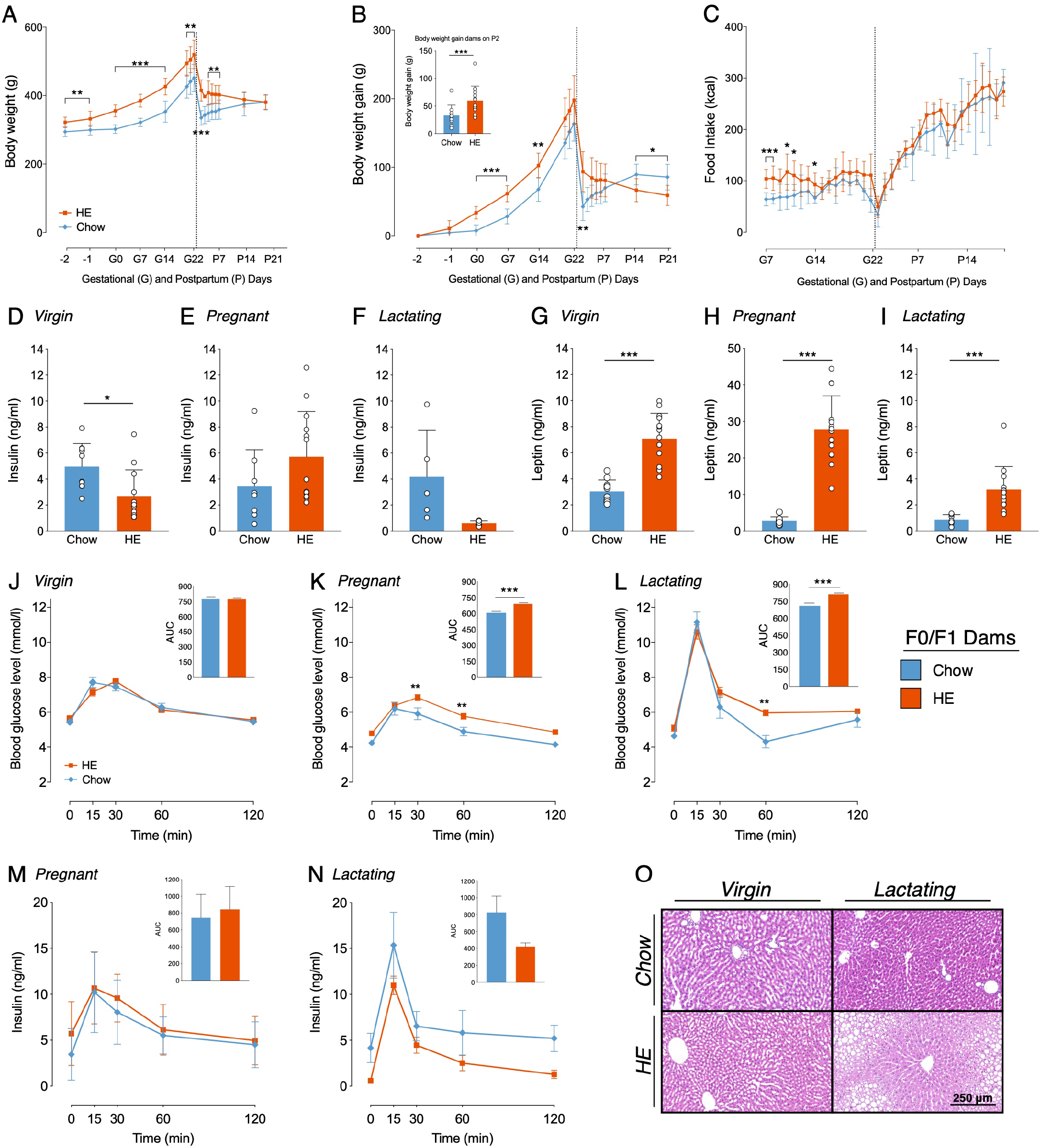
Influence of HE-diet on physiological parameters in obesity-prone and - resistant rat dams in filial generation 0 and 1 (F0/F1) before, during and after pregnancy. Body weight in grams (A), body weight gain in grams (B) and food intake in kilocalories (C) of Sprague Dawley dams on chow (blue) or high energy (HE, orange) diet over the course of pregnancy and lactation. Vertical dotted line marks parturition. (B) Inset shows body weight gain during pregnancy (difference between body weight on gestational day 1 (G1) and P2 to account for litter and placenta weight). (D-I) Plasma insulin and leptin levels (ng/ml) following a 12h fast and prior to an oral glucose tolerance test (OGTT) in dams on chow and HE diet. (D, G) Insulin and leptin levels in virgin dams following a one week HE challenge. (E, H) Insulin and leptin levels during mid-pregnancy (gestational day 15) in dams on chow or HE diet. (F, I) Insulin and leptin levels in mid-lactation (postpartum day 12) in dams on chow or HE diet. (J-L) Blood glucose levels (mmol/l) at different time points (0, 15, 30, 60, 120 min) during an oral glucose tolerance test in virgin (J), pregnant (K) and lactating (L) dams on chow or HE diet. Inset in each graph J-L shows the area under the curve relative to zero. (M-N) Insulin levels (ng/ml) during the OGTT in pregnant (M) and lactating (N) rats on either chow or HE diet. Insets show area under the curve relative to 0 ng/ml. (O) Representative HE-stained liver sections of virgin and lactating dams on either chow or HE diet (Scale Bar: 250 μm). Data are represented as mean ± SD. (A-C) 2-way-ANOVA respective mixed effect model to account for missing values with Sidak’s multiple comparisons post-hoc test. (D-I) two-tailed Welch’s test. J-N 2-way-ANOVA with Sidak’s multiple comparisons post-hoc test. Insets (J-N) two-tailed unipaired t-tests. *p < 0.05, **p < 0.01, *** p < 0.001. N = 10 in chow group, N = 14 in HE group.

#### Leptin and Insulin levels

Insulin and leptin were measured in plasma following a 12-h fast and prior to an OGTT performed before mating (1 week following the HE-challenge), during mid-pregnancy (G15), and mid-lactation (P12). Prior to mating, insulin values were lower in the prospective HE-fed females than those maintained on chow (t_20_ = 2.774, P = 0.012; Fig. 2D], but were not different during pregnancy (Fig. 2E) or lactation (Fig. 2F). Leptin, however, was significantly elevated in the HE-fed females prior to (t_19.22_ = 6.851, P < 0.001; Fig. 2G), during (t_13.69_ = 10.14, P < 0.001; Fig. 2H), and after pregnancy (t_14.03_ = 4.526, P < 0.001, Fig. 2I).

#### Oral glucose tolerance test

Intake of HE also influenced glucose clearance during an OGTT in pregnant and lactating, but not virgin, rat dams. In virgin rats a main effect of time (F_4,88_ = 99.3, P < 0.001) was found as well as a significant interaction between time and diet (F_4,88_ = 2.886, P = 0.027; Fig. 2J]. In pregnant rats (Fig. 2K), main effects of diet (F_1,21_ = 24.58, P < 0.001) and time (F_4,84_ = 45.62, P < 0.001) were detected, with HE-fed rats exhibiting higher blood glucose levels than chow-fed rats at 30 min (t_105_ = 3.351, P = 0.006) and 60 min (t_105_ = 3.2, P = 0.009) after the glucose challenge. During lactation (Fig. 2L), main effects of diet (F_1,21_ = 7.414, P = 0.013) and time (F_4,84_ = 98.29, P < 0.001) were observed, as well as a significant interaction (F_4,84_ = 2.627, P = 0.04). Significantly higher blood glucose levels in HE-fed dams compared to chow-fed were found at the 60 min time point (t_105_ =3.389, P = 0.005). AUC analysis, relative to zero, demonstrated significantly higher AUCs in pregnant and lactating HE-fed rats compared to pregnant or lactating chow-fed rats, respectively (Pregnancy t_21_ = 5.176, P < 0.001, Lactation t_21_ = 4.3, P < 0.001; Fig. 2K and 2L insets).

Insulin levels during the OGTTs performed in pregnant and lactating dams showed a similar curve pattern as blood glucose levels, with a peak in plasma insulin concentration 15 min after the glucose challenge (Figs. 2M and 2N). In pregnant rats, only a main effect of time (F_4,76_ = 25.4, P < 0.001) was found, whereas in lactating rats main effects of diet (F_1,11_ = 8.592, P = 0.014) and time (F_4,44_ = 23.34, P < 0.001) were detected. However, no significant differences in plasma insulin concentrations were found comparing pregnant or lactating chow-fed rats to pregnant or lactating HE-fed rats. AUC, relative to 0 ng/ml, did not show any significant differences in insulin concentration between chow- and HE-fed rats.

#### Liver Histology

We observed no influence of HE diet following histological inspection of visceral adipose tissue and pancreas collected from F0/F1 dams. In the livers, however, we observed mild to severe lipidosis in all livers from HE-fed lactating dams (average score was 1.93 ± 0.83), while all chow-fed dams received a score of 0, indicating no lipidosis. What was perhaps most interesting, was the observation that three female rats that were maintained on HE diet for the same duration as the dams, but who did not become pregnant, also all received a score of 0, suggesting that liver lipidosis resulted from an interaction between HE consumption and the states of pregnancy and lactation. Representative images from virgin or lactating females, fed chow or HE, are shown in Figure 2O.

#### Meal patterns

Analysis of meal size (Fig. 3A) revealed a main effect of diet (F_1,97_ = 113.4, P < 0.001) and time (F_4,97_ = 26.22, P < 0.001) as well as significant interaction (F_4,97_ = 3.183, P = 0.018). Meal size was significantly larger in HE-fed rats than in chow-fed rats for all pregnant and postpartum measurement days: G7 (t_97_ = 4.305, P < 0.001), G20 ± 1 (t_97_ = 2.905, P = 0.024), P4 (t_97_ = 4.449, P < 0.001), P9 (t_97_ = 5.696, P < 0.001) and P15 ± 1 (t_97_ = 7.659, P < 0.001).

**Figure 3.**
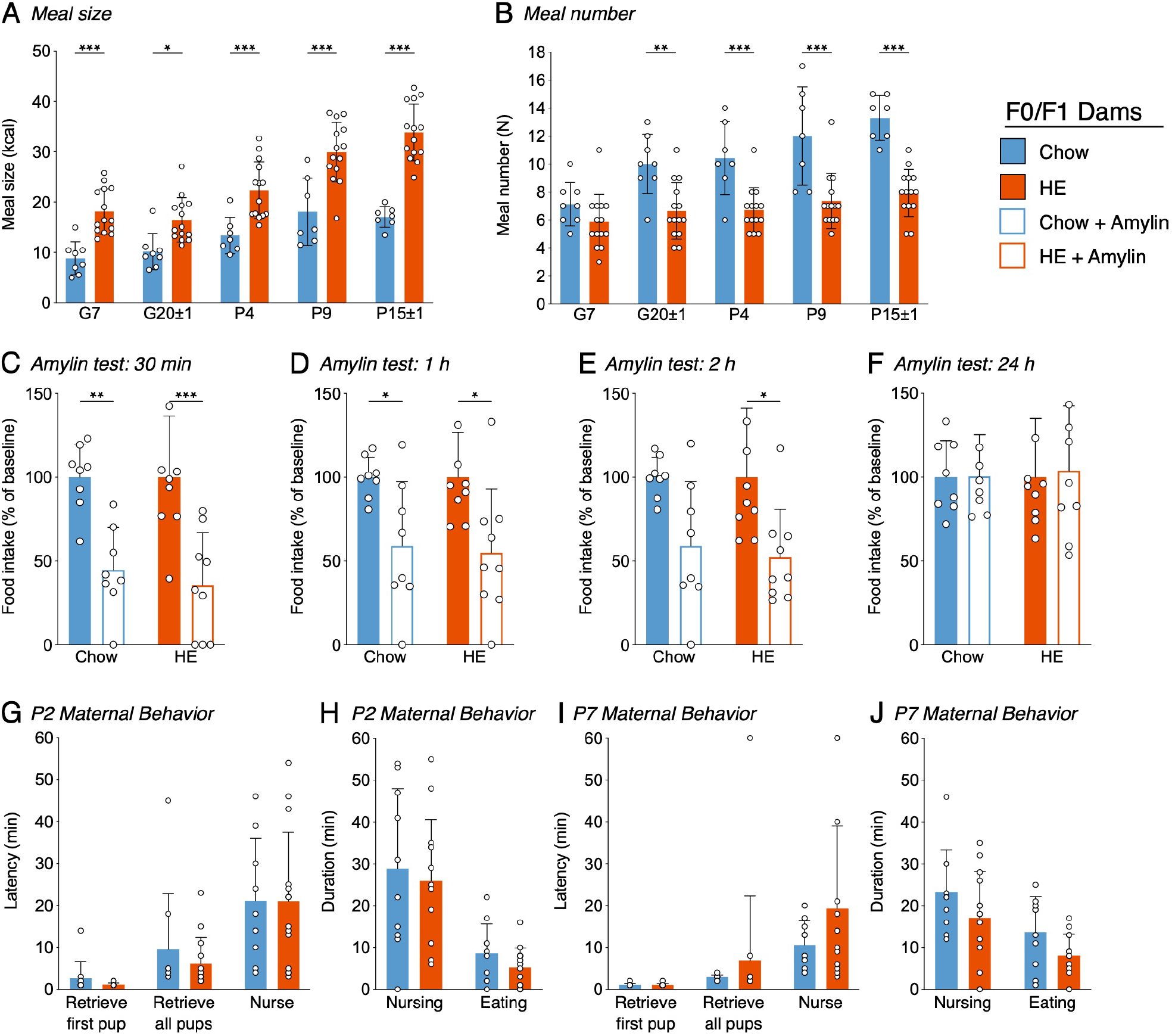
Influence of HE-diet in obesity-prone and -resistant rat dams in filial generation 0 and 1 (F0/F1) on meal pattern, amylin sensitivity and maternal behavior. Meal size in kilocalories (A) and meal number (B) of pregnant and postpartum rat dams on chow or high energy (HE) diet on gestational day 7 (G7), G20±1, postpartum day 4 (P4), P9, and P15±1. (C-F) Food intake as percentage of baseline following a single dose subcutaneous amylin (20 μg/kg) or vehicle (saline) injection after a 4-h fast in rat dams on gestational day 11±2, 30 minutes (C), 1 hour (D), 2 hours (E) and 24 hours (F) after injection. (G-J) Pup retrieval test and 60 minutes maternal behavior home cage observation in dams on either chow or HE diet on postpartum day 2 (P2) and 7 (P7) in the first four hours after dark onset. Latency to retrieve first pup, latency to retrieve all pups and latency to start nursing in minutes on P2 (G) and P7 (I). Total nursing and eating duration in minutes on P2 (I) and P7 (J). Data are represented as mean ± SD. (A-F) 2-way-ANOVA respective mixed effect model to account for missing values with Sidak’s multiple comparisons post-hoc test; (G-J) two-tailed unpaired t-tests. *p < 0.05, **p < 0.01, *** p < 0.001. N = 10 in chow group, N = 14 in HE group.

Meal number (Fig. 3B) was significantly lower in HE-fed rats compared to chow-fed rats. Main effects of diet (F_1, 97_ = 77.98, P < 0.001) time (F_4, 97_ = 11.31, P < 0.001), and an interaction (F_4, 97_ = 2.85, P = 0.028) were detected. Meal number was significantly decreased in HE-fed rats compared to chow-fed rats on G20 ± 1 (t_97_ = 1.40, P = 0.002), P4 (t_97_ = 3.71, P < 0.001), P9 (t_97_ = 4.92, P < 0.001) and P15 ± 1 (t_97_ = 5.67, P < 0.001).

#### Amylin sensitivity test

Because pregnancy causes reduced sensitivity to hormones regulating food intake, including leptin and cholecystokinin [19; 38], we assessed if the action of the satiating hormone amylin is preserved in pregnant rat dams maintained on either chow or HE diet (Fig. 3C-F). By representing the food intake following amylin-treatment as a percentage of baseline (vehicle food intake = 100%), the effectiveness of amylin was compared across diet groups on pregnancy day G11 ± 3. Overall, amylin’s effectiveness to reduce food intake was similar between diet groups, and consistent with what is observed in non-pregnant rats [39]. There was a main effect of amylin treatment on food intake at 30 min (F1,15 = 31.82, P < 0.001; Fig 3C), 1 h (F1,15 = 14.76, P = 0.002; Fig. 3D), and 2 h (F1,15 = 13.29, P = 0.02; Fig. 3E), but never an effect of diet. By 24 h, all treatment effects were gone (Fig. 3F).

#### Maternal behavior

A pup retrieval test followed by 60 min of home-cage observations of maternal behaviors was conducted on P2 and P7 in dams fed chow or HE. We observed no effect of maternal diet on the latency to retrieve pups to the nest or begin nursing on P2 (Fig. 3G) or P7 (Fig. 3I). When using the durations of nursing behavior and eating behavior to approximate pup- and dam-related behavior, we again observed no effect of diet on these parameters on P2 (Fig. 3H) or P7 (Fig. 3J).

### Experiment Two: Influence of HE and polygenic predisposition on postpartum metabolic and behavioral outcomes in F3 dams

#### Maternal body weight gain and food intake

An overall effect of time (F_3.708, 91.41_ = 316.1, P < 0.0001), phenotype (F_1, 25_ = 23.56, P < 0.0001), time x phenotype interaction (F_29, 715_ = 3.549, P < 0.0001) and time x diet interaction (F_29, 715_ = 18.80, P < 0.0001) on body weight was found (Fig. 4A). DR chow-fed rats had significantly lower body weight compared to DIO chow-fed animals already starting from day G1 until the end of the experiment on P25 (average P < 0.001). Furthermore, DR chow-fed animals were significantly lighter than DIO HE-fed animals at P1 (P = 0.04). DR HE-fed rats were significantly lighter than DIO chow-fed animals from day P6-P22 (average P = 0.01). In addition, DR-chow animals dropped more body weight from P1 to P3 compared to DR HE-fed dams (average P = 0.002).

**Figure 4.**
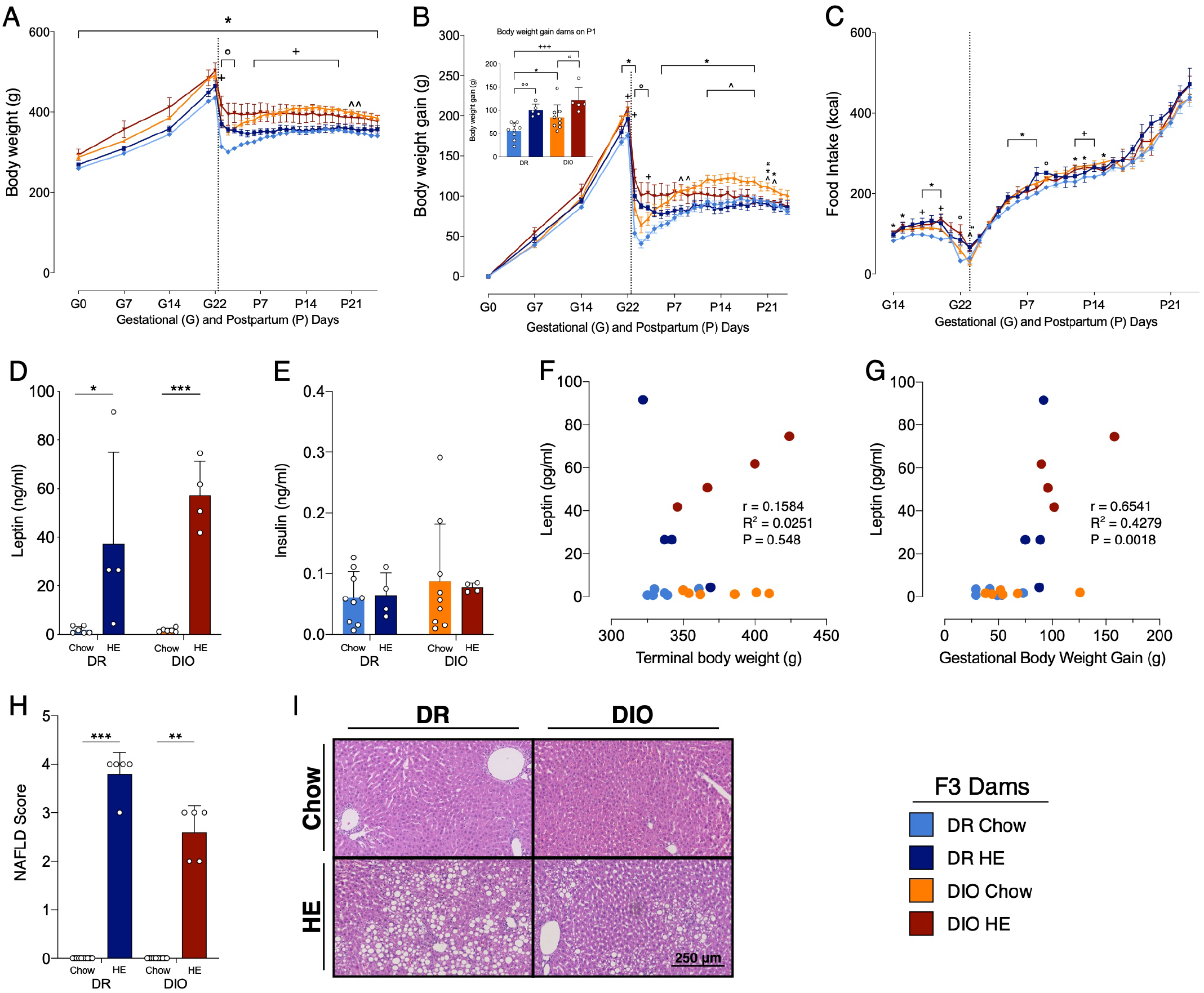
Influence of HE-diet on physiological parameters in diet resistant (DR) and diet induced obese (DIO) rat dams in filial generation 3 (F3) before, during and after pregnancy. Body weight in grams (A), body weight gain in grams (B) and food intake in kilocalories (C) of DR dams on chow (light blue), or HE diet (dark blue), DIO dams on chow (orange), or HE diet (red) over the course of pregnancy and lactation. Vertical dotted line marks parturition. (B) Inset shows body weight gain during pregnancy (difference between body weight on gestational day 1 (G1) and postpartum day 1 (P1) to account for litter and placenta weight). (D-E) Leptin (D) and insulin (E) levels (ng/ml) in plasma of DR and DIO dams on chow or HE diet at sacrifice around postpartum day 21±1. Animals on HE displayed higher plasma leptin levels compared to dams on chow. (F-G) Scatter plot of terminal leptin levels (pg/ml) and terminal body weight (g) (F) or gestational body weight gain (g) (G) in DR and DIO dams on chow or HE diet. Terminal leptin levels and gestational body weight gain correlate. (H) Severity of liver lipidosis according to the NAFLD scoring system for rodents (Grade 0-4) in DR and DIO dams on chow or HE diet. HE diet triggered liver lipidosis regardless of phenotype. Representative images of H&E stained liver sections in Fgure I. Data are represented as mean ± SD. (A-E) 2-way-ANOVA respective mixed effect model to account for missing values with Tukey’s multiple comparisons post-hoc test. (F-G) Correlation Pearson r statistical test. (H) Kruskal-Wallis test with Dunn’s post hoc test *p < 0.05, **p < 0.01, *** p < 0.001. DR-chow (N = 10), DR-HE (N = 5), DIO-chow (N = 9), and DIO-HE (N = 5).

Similar effects were observed for the body weight gain (Fig. 4B). There was an overall effect of time (F_3.294, 81.99_ = 332.6, P < 0.0001), phenotype (F_1, 25_ = 12.14, P = 0.0018), a time x phenotype interaction (F_28, 697_ = 3.348, P < 0.0001) and a time x diet interaction (F_28, 697_ = 19.50, P < 0.0001). DR chow-fed dams had significantly less body weight gain compared to DIO chow-fed animals from G21-P1 and from P5 to P22 (average P = 0.01). DR chow-fed animals showed reduced body weight gain compared to DIO HE-fed animals from G22 to P3 (average P = 0.03). From P12 up to P19 DIO chow-fed animals gained significantly more weight compared with DR HE-fed animals (average P = 0.02). Comparing body weight gain, chow-fed animals had a greater body weight gain from P1 to P3 in the DR group and from G22 to P3 in the DIO group compared with their HE-fed counterparts, with a stronger effect in the DR strain (DR-Chow vs. DR-HE average P < 0.001; DIO-Chow vs. DIO-HE average P = 0.03).

Comparison of body weight gain during pregnancy (the difference between body weight on G1 and on P1 to account for litter and placenta weight, See inset in Fig. 4B) revealed a phenotype effect (F_1, 24_ = 8.125, P = 0.009) and diet effect (F_1, 24_ = 21.04, P < 0.001), without a significant interaction (F_1, 24_ = 0.2153, P = 0.65). Multiple comparison analysis showed that DR chow-fed animals gained less body weight during pregnancy compared to DR HE-fed dams (P = 0.008), as well as compared to DIO chow-fed dams (P = 0.05). Further, DIO HE-animals gained more body weigh compared to DIO chow-fed dams (P = 0.04). The biggest difference in weight gain could be observed between DR chow-fed dams and DIO HE-fed animals (P < 0.001). DIO HE-fed gained 119% more gestational weight during pregnancy compared to the DR chow-fed group.

When food intake was corrected for the energy density per gram food in kilocalories (kcal) there was a time effect (F_7.143, 167.5_ = 431.0, P < 0.0001), a diet effect (F_1, 24_ = 16.98, P = 0.0004), a time x diet interaction (F_31, 727_ = 1.471, P = 0.0489) and a phenotype x diet interaction (F_1, 24_ = 6.380, P = 0.0185; Fig. 4C). Multiple comparisons detected only few significant differences between groups on specific days. DR chow-fed dams ate significantly less kcal than DR HE-fed animals on day G22 and P9 (average P = 0.021). Further, DIO chow-fed rats ate significantly more kcal on G15-G16, G18-G20 and from P5 to P8 compared to DR chow-fed dams, showing a tendency for DIO animals to eat more energy during gestation and lactation (average P = 0.015). In addition, chow-fed animals exhibited a more pronounced decrease in kcal intake around P1 compared to HE-fed animals (DR-Chow vs. DR-HE on G22 P = 0.04; DIO-Chow vs. DIO-HE on P1 P = 0.01; DR-HE vs. DIO-Chow on P1 P = 0.02). On G18 and G20 and from P12 to P14 DR chow-fed animals ate significantly less kcal compared to DIO HE-fed animals (average P = 0.36).

#### Postpartum leptin and Insulin levels

Postpartum leptin and insulin levels were measured at day of sacrifice around postpartum day 21 ± 1. Analysis of leptin levels revealed a diet effect (F_1, 16_ = 32.52, P = < 0.0001) with no phenotype effect (F_1, 16_ = 1.546, P = 0.2316) and no interaction (F_1, 16_ = 1.586, P = 0.2260, Fig. 4D). Multiple comparisons showed significant differences between DR chow and DR HE (P = 0.0289), DR chow vs. DIO HE (p = 0.0008), DR HE vs. DIO chow (p = 0.0283) and DIO chow vs. DIO HE (p = 0.0008), with animals fed a HE-diet having higher leptin levels than animals on chow. There was no significant difference in insulin levels between diet or phenotype groups (Fig. 4E). Further, no significant correlation was found when correlating leptin levels to terminal body weight (Fig. 4F). Interestingly a correlation between blood leptin levels and gestational body weight gain could be observed (r = 0.6541, R^2^ = 0.4279, p = 0.0018, Fig. 4G).

#### Histological analysis of postpartum liver

Consistent with observations in the first experiment, there was mild to severe lipidosis in all livers from HE-fed lactating dams (average score DR: 3.8 ± 0.45; DIO: 2.6 ± 0.55), while all chow-fed dams displayed no liver lipidosis and received a score of zero. Significant differences between groups could be detected using a Kruskal-Wallis test (p < 0.001). Dunn’s multiple comparison revealed significant differences between DR chow vs. DR HE (p < 0.001) and DIO chow vs. DIO HE (p = 0.006) groups (Fig. 4H). Representative H&E-stained livers from each group are shown in Figure 4I.

#### Meal pattern analysis

Analysis of meal size and meal number revealed the same tendencies as described in generations F0/F1. For meal size (Fig. 5A), we detected a main effect of time (F_2.617, 64.55_ = 65.55, P < 0.001), diet (F_1, 25_ = 120.3, p < 0.001), time x diet interaction (F_3, 74_ = 14.68, p < 0.001) and time x phenotype x diet interaction (F_3, 74_ = 3.760, p = 0.0143), with animals on HE diet having larger meal size compared to animals on chow, regardless of phenotype. Meal number (Fig. 5B) was significantly lower in HE-fed rats compared to chow-fed rats, again regardless of phenotype. Main effects of time (F_2.680, 65.21_ = 13.86, p < 0.001), diet (F_1, 25_ = 64.88, p < 0.001), and time x diet interaction (F_3, 73_ = 4.894, p = 0.0037) were detected.

**Figure 5.**
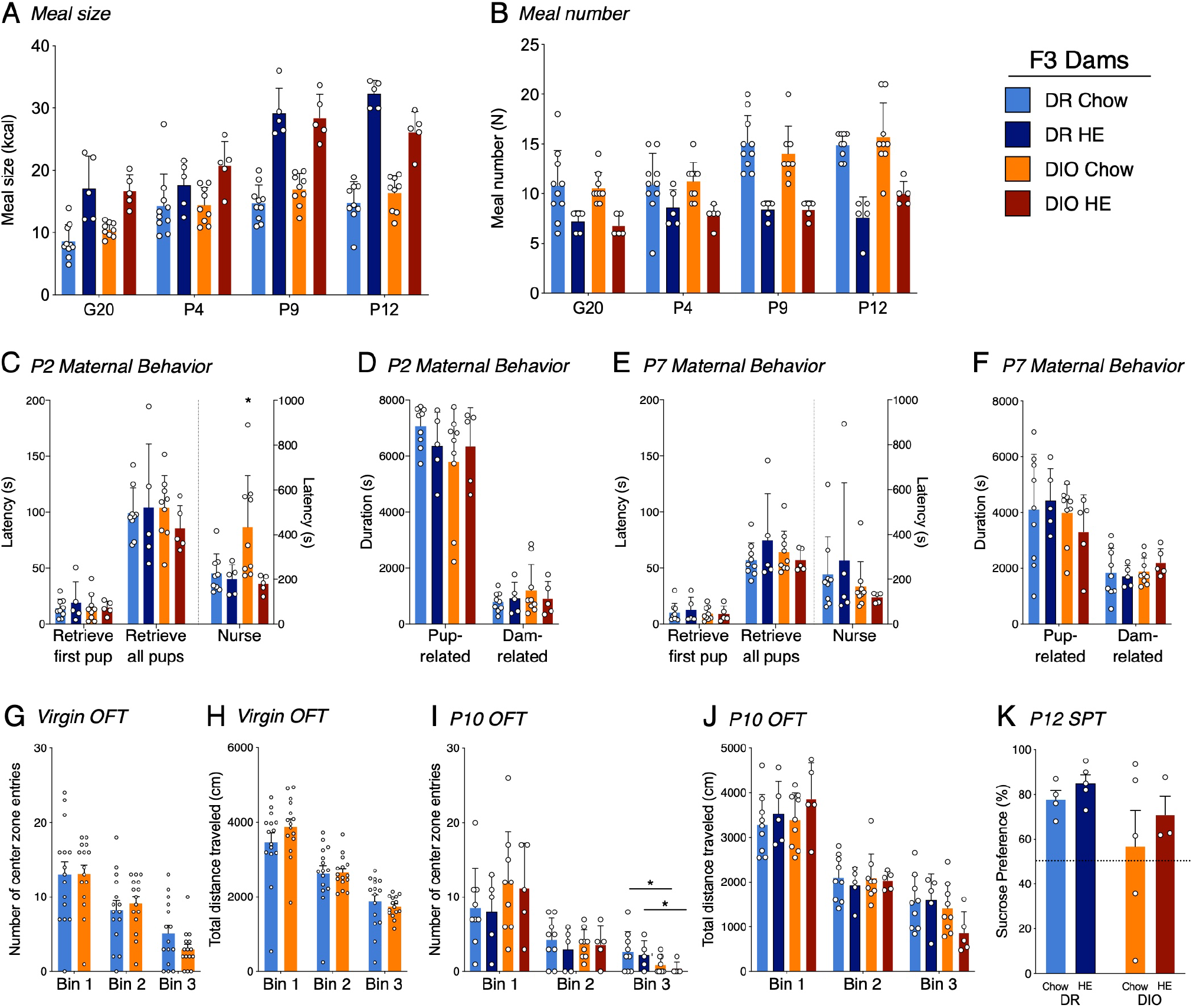
Influence of HE-diet in diet resistant (DR) and diet induced obese (DIO) rat dams in filial generation 3 (F3) on meal pattern, amylin sensitivity and maternal behavior. Meal size in kilocalories (A) and Meal number (B) of pregnant and postpartum DR and DIO rat dams on chow or high energy (HE) diet on gestational day 20 (G20), postpartum day 4 (P4), P9, and P12. (C-F) Maternal behavior in DR and DIO dams on chow or HE diet on postpartum day 2 (P2) or 7 (P7). Latency to retrieve first pup, latency to retrieve all pups and latency to start nursing in seconds on P2 (C) and P7 (E). Total duration in seconds of pup related and dam related behavior on P2 (D) and P7 (F). (G-H) Open field test in virgin DR and DIO rat dams around 9 weeks of age on chow. (G) Number of center zone entries divided into 3 bins, whereas one bin represents 5 minutes of tested time. (H) Total travelled distance in centimeter (cm) in virgin DR and DIO dams per bin. (I-J) Open field test on postpartum day 10 in DR and DIO rat dams on chow or HE diet. Number of center zone entries (I) and total traveled distance in centimeter (J) divided into 3 bins, representing 5 minutes of test each. (K) Sucrose preference in percent of total liquid intake on postpartum day 14-15 in DR and DIO rat dams on either chow or HE diet. 1% sucrose solution and water intake were averaged over two consecutive days. Animals displaying a clear side preference were excluded. Data are represented as mean ± SD (A-K) 2-way-ANOVA respective mixed effect model to account for missing values with Tukey’s multiple comparisons post-hoc test. *p < 0.05, **p < 0.01, *** p < 0.001. DR-chow (N = 10), DR-HE (N = 5), DIO-chow (N = 9), and DIO-HE (N = 5).

#### Maternal behaviors

On P2, while there were no differences between the groups for latency to retrieve pups to the nest, DIO-chow dams demonstrated an increased latency to begin nursing their pups (P = 0.0023; Fig. 5C), which was significantly higher than all other groups. On P7, though there were no differences between latencies to retrieve or nurse (Fig. 5E), or the total duration of pup- or dam-related behaviors across groups (Fig. 5F), rats on HE diet showed an increased time dedicated to pup grooming compared with chow-fed rats (F_1, 24_ = 5.281, P = 0.03), whereas phenotype had no effect (F_1, 24_ = 0.02550, P = 0.87; See Supplemental Results). On all the other behaviors, and regardless of postpartum day 2 or 7, there were no further statistical differences between the different groups (See Supplemental Results for details). For most behaviors measured, neither phenotype nor diet influenced the maternal behavior on P2 or P7.

#### Open Field Test

To assess the physical activity and anxiety level, rat dams were tested in an open field test prior to pregnancy and again at postpartum day 10 or 11. Latency to enter the center zone, percentage of time spent in the center zone, total distance per 5-minute bin, as well as center zone distance and center zone entries per bin were measured. To compare baseline results unpaired t-tests were used (Fig. 5G and 5H). There were no statistical differences between DIO and DR animals in any of the assessed parameters. At the time point of baseline measurements, all dams were maintained on chow diet, hence a diet effect could not be measured.

To compare the different parameters on postpartum day 10 or 11, a two-way ANOVA with Tukey’s multiple comparisons test was used (Fig. 5I and 5J). A phenotype effect was observed for the number of center zone entries in Bin 3; DIO rat dams entered the center zone fewer times in the last 5 minutes of the test, regardless of the diet (F_1, 24_ = 5.682, P =0.03; Fig. 5I). In all other measurements there were no significant differences between phenotype nor diet.

#### Sucrose Preference Test

A sucrose preference test was performed on postpartum day 16. Rats demonstrating a clear side-preference (i.e. consistently drinking from the left or right bottle regardless of content) were excluded from the analysis. There was neither a diet effect (F1, 13 = 0.9778, P = 0.34) nor a phenotype effect (F1, 13 = 2.667, P = 0.13; Fig. 5K) detected.

### Experiment Three: Influence of HE and polygenic predisposition on select postpartum outcomes in F6 and F7 dams

#### Maternal body weight gain

Select parameters were measured in later generations of the DR and DIO dams. Most data in the following results were collected from F7 dams, however the OGTT, plasma leptin, and SPT also included data from F6 dams. In line with what was observed in the F3 dams, analysis of body weight gain data (Fig. 6A) in pregnancy and postpartum revealed main effects of phenotype (F_1, 15_ = 7.007, P = 0.018), diet (F_1, 15_ = 9.093, P = 0.009), and time (F_2.616, 38.23_ = 427.0, P < 0.001). Individual group differences detected during late pregnancy were mainly a phenotype effect, meaning that DIO dams gain more weight during pregnancy. The group differences detected in early postpartum were mainly a diet effect, reflecting that chow-fed dams have a more pronounced body weight drop after parturition.

**Figure 6.**
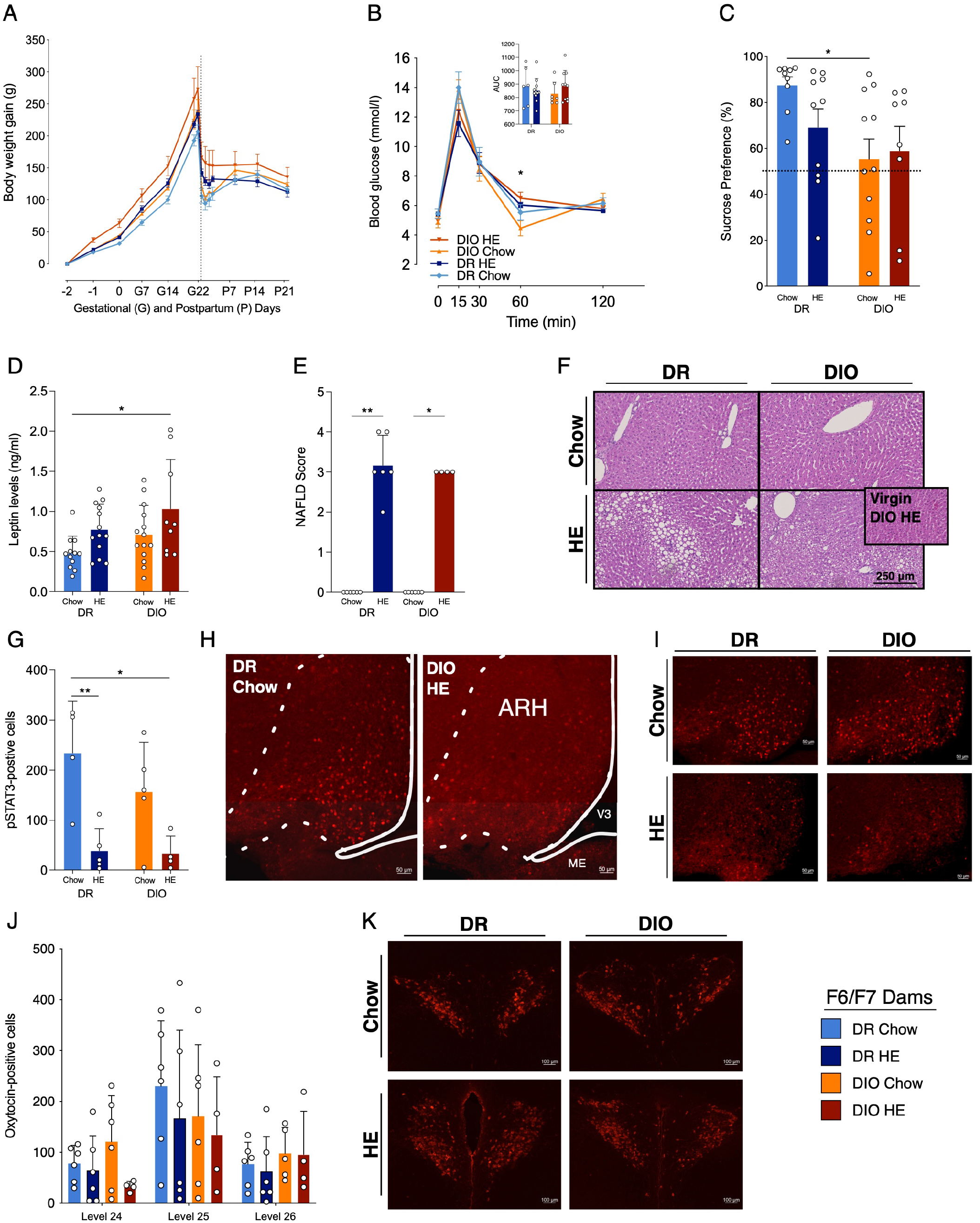
Influence of HE-diet in diet resistant (DR) and diet induced obese (DIO) rat dams in filial generation 6 and 7 (F6, F7) on selected physiological parameters. (A) Body weight gain in gram (g) of DR dams on chow (light blue), or HE diet (dark blue), DIO dams on chow (orange), or HE diet (red) over the course of pregnancy and lactation. Dotted vertical line marks partuition. (B) Blood glucose levels (mmol/l) at different time points (0, 15, 30, 60, 120 min) during an oral glucose tolerance test in DR and DIO dams on chow or HE diet. Inset shows the area under the curve (AUC) relative to zero. (C) Sucrose preference in percent of total liquid intake on postpartum day 14-15 in DR and DIO rat dams on either chow or HE diet. 1% sucrose solution and water intake were averaged over two consecutive days. Animals displaying a clear side preference were excluded. (D) Plasma leptin levels in ng/ml in DR and DIO dams on chow or HE diet after a 12-h fast on P18±1. Dams on HE displayed a higher leptin concentration compared to dams on chow. (E) Severity of liver lipidosis according to the NAFLD scoring system for rodents (Grade 0-4) in DR and DIO dams on chow or HE diet. HE diet triggered liver lipidosis regardless of phenotype. (F) Representative images of H&E sections. Inset in (F) shows image from a nonpregnant, non-lactating DIO dam after 6 weeks on HE diet (no lipidosis present). (G) Immunohistochemistry of pSTAT3-positive cells in the arcuate nucleus of the hypothalamus (ARH) in DR and DIO dams on chow or HE diet after 5 mg/kg intraperitoneal leptin injection forty-five minutes prior to sacrifice on postpartum day 21±1. Animals on chow showed greater leptin-induced pSTAT3 phosphorylation compared to animals on HE diet. (H) Overview of the ARH in a DR dam on chow diet and a DIO dam on HE diet. (I) Zoomed in representative pictures of the ARH in each group. (J) Oxytocin positive cells in the paraventricular nucleus of the hypothalamus (PVH) on different brain levels (level 24-26) (according to Swanson brain atlas) in DR and DIO dams on chow or HE diet. (K) Representative pictures of the PVH on level 25 in each group. Data are represented as mean ± SD (A-D, G, J) 2-way-ANOVA respective mixed effect model to account for missing values with Tukey’s multiple comparisons post-hoc test. (E) Kruskal-Wallis test with Dunn’s post hoc test. *p < 0.05, **p < 0.01, *** p < 0.001. DR-chow (F6 N = 8, F7 N = 6), DR-HE (F6 N = 7, F7 N = 6), DIO-chow (F6 N = 8, F7 N = 6), and DIO-HE (F6 N = 6, F7 N = 4).

#### Oral Glucose Tolerance Test

OGTT were conducted in F6 and F7 dams on P18 (Fig. 6B). Of note, blood glucose was measured before and after administration of the glucose load via gavage, unlike in F0/F1 dams which consumed the glucose orally. A mixed-effect model analysis of the data revealed a main effect of time (F_2.206, 68.40_ = 142.6, P < 0.001), and an interaction between time and diet (F_4, 124_ = 4.645, P = 0.002). While there were no differences in the AUC of the four groups, an individual difference between DIO-Chow and DIO-HE dams was detected at 60 min (P = 0.036) with Tukey’s multiple comparisons test.

#### Sucrose Preference Test

While the data from the SPT in F3 dams suggested a trend for DIO dams to exhibit reduced preference for sucrose, a significant main effect of phenotype was observed in the F6/F7 dams (F_1, 34_ = 6.604, P = 0.015), with Tukey’s multiple comparisons test revealing an individual difference in sucrose preference between the DR-Chow and DIO-Chow dams (P = 0.037; Fig. 6C).

#### Postpartum leptin levels

Following analysis of plasma leptin levels collected at timepoint 0 during the OGTT on P18 (Fig. 6D), main effects of phenotype (F_1, 44_ = 4.891, P = 0.032) and diet: (F_1, 44_ = 7.571, P = 0.009) were detected, with Tukey’s multiple comparisons test revealing a significant difference between the leptin levels in DR-Chow versus DIO-HE dams (P = 0.010).

#### Histological analysis of postpartum liver

The presence and extent of lipidosis was again assessed in F7 dams, and the results were consistent with the findings from F3 dams. The semi-quantitative scoring (Kruskal-Wallis: P < 0.001; Fig. 6E) again detected difference between DR-chow and DR-HE (P = 0.003) and DIO-chow and DIO-HE (P = 0.211) livers. Representative liver images from each group are shown in Fig. 6F. Note that the inset in the DIO-HE image is from a DIO female that failed to become pregnant, but was maintained on HE for the same duration as the DIO-HE dams; this nonlactating case received a score of 0.

#### Postpartum leptin-induced pSTAT3 in ARH

Analysis of the number of leptin-induced pSTAT3-positive neurons in the arcuate nucleus of the hypothalamus (ARH; Fig. 6G) in DR and DIO dams fed chow or HE diet, detected a main effect of diet (F_1, 14_ = 19.13, P < 0.001). Multiple comparisons using Tukey’ post hoc test, showed that DR-chow dams have more pSTAT3-positive neurons compared to DR-HE (p = 0.010), as do DR-chow compared to DIO-HE dams (p = 0.011). In Figures 6H and 6I, representative images of pSTAT3-positive neurons in the ARH (Level 29 in [35]) are shown.

#### Postpartum levels of oxytocin in PVH

Number of oxytocin-positive cells were evaluated following immunohistochemical staining in the paraventricular nucleus of the hypothalamus (PVH). Figure 6J shows the number of oxytocin-positive neurons in the three brain levels quantified (Level 24-26 in [35]). Using a three-way ANOVA, we could detect significant differences across different brain levels (F_2, 53_ = 7.022, P = 0.002), however, no further statistical differences could be observed across groups in the PVH at this point of late lactation. In Figure 6K, representative images of oxytocin-positive neurons from brain Level 25 (in [35]) are shown.

## Discussion

More than one third of women of reproductive age in the US are obese [1]. This prevalence of obesity, driven in part by an overconsumption of a palatable, high energy diet and a polygenic-sensitivity to such diets, continues to increase [40; 41], further propagating the negative health consequences of maternal obesity for mother and child. While many studies in humans and rodents have investigated the impact of obesity and an unhealthy diet during pregnancy on the offspring, few studies have evaluated how this metabolic challenge effects the physiological parameters of the mother, despite the evidence that women with obesity are at higher risk for adverse health effects during and after pregnancy, such as gestational diabetes and postpartum depression [11; 42].

The aim of this project was to characterize the physiological status of a selectively bred diet-induced obese (DIO) and diet-resistant (DR) rat model fed a high energy diet (HE) or normal chow in a novel context of the metabolically demanding periods of pregnancy and lactation. To achieve this goal, we monitored physiological parameters such as maternal body weight, body weight gain, food intake, blood leptin and insulin levels and oral glucose tolerance in F0/F1 generations, for which the HE diet was the main factor. The offspring from these experiments were selectively bred according to the protocol developed by Levin and colleagues, whereby the highest and lowest weight gainers were bred to generate DIO and DR rat lines [24]. In a second project we examined the influence of diet and phenotype on maternal behaviors in the F3 generation, performing high-resolution tests of maternal behaviors, open field tests to determine activity and anxiety-like behaviors, and sucrose preference tests as a readout for hedonic behavior, while maintaining close monitoring of body weight, body weight gain and food intake of the dams and corresponding litters. In the third phase of the study, in addition to monitoring select physiological and behavioral parameters, dams from generations F6 and F7 were used to assess leptin sensitivity in the postpartum brain, and whether these factors of diet and genetic proneness for obesity might alter expression of oxytocin neurons in the paraventricular nucleus of the hypothalamus. Collectively, our studies show that while consumption of HE diet during and after pregnancy can influence maternal physiology and behavior, selective breeding based on the sensitivity to this diet can have both independent and compounded effects on maternal health.

### Effects of diet and selective breeding on gestational body weight and leptin levels

To assess the effect of phenotype and diet on physiological and metabolic parameters body weight, body weight gain, food intake, leptin and insulin levels were monitored throughout the different generations. We could show that by selectively breeding DIO and DR rat strains, phenotype differences regarding body weight and body weight gain can be enhanced, starting from generation F3, with the DIO line having elevated BW and BWG compared to DR dams of the same age regardless of diet. When comparing dams on chow with dams on HE diet, an overall diet effect is already visible starting in the F0/F1 generation, with HE animals having increased BW and BWG compared to chow dams. The biggest difference was observed between DR chow-fed and DIO HE-fed dams in generation F3, with DIO HE-fed animals gaining 119% more gestational weight from G0 to P1 during pregnancy. Following parturition, dams on chow diet displayed a dramatic drop in body weight, which was followed by a slow but steady regain of body weight mainly from P2 to P12. In contrast, HE dams stayed nearly stable with their body weight after parturition until P25. This effect of HE on the pattern of postpartum body weight changes was observed over several generations, and a similar trend can be observed in recently published work in mouse dams [15]. While the significance of this pattern will require additional probing, the consistency of the effect suggests that intake of HE can influence the metabolic adaptions that occur from pregnancy to postpartum. Consistent with the elevated gestational weight gain, dams fed HE had increased leptin levels during and after pregnancy compared to chow-fed dams. One of our central hypotheses is that excess adiposity or dietary fat during and after pregnancy delays or prevents the return of leptin sensitivity in the postpartum period, to which high maternal levels of circulating leptin likely contribute.

### Effects of diet and selective breeding on leptin resistance

Leptin resistance is a known and necessary metabolic adaptation that enables maternal fat accumulation during pregnancy [18; 19; 20], providing energy to the growing fetus and suckling newborn. The progression of gestational leptin resistance has been documented, and while its reversal after pregnancy is assumed [43], the time course of its reversal has not been traced. Here we provide initial evidence that intake of HE diet during pregnancy and lactation blunts hypothalamic leptin sensitivity, as demonstrated by reduced leptin-induced pSTAT3 in the brains of HE-fed dams at the time of pup-weaning (P26). We observed this effect of HE in both DR and DIO dams. The precise consequence of retained postpartum leptin resistance has also not been investigated, but could be far-reaching provided the strong connections between metabolic sensing and signaling in the brain and various other brain pathways controlling motivation and cognition [44; 45], let alone the impact of leptin signaling in the periphery on energy homeostasis [46]. For instance, deficits in postpartum leptin function as a result of HFD-intake in mice was recently identified as contributor to insufficient prolactin signaling and lactation [47]. And in women, elevated early postpartum leptin levels, which likely coincide with retained leptin resistance, was shown to be predictive of postpartum depression [48].

Leptin resistance during pregnancy and in the postpartum period might underlie other consequences of HE intake in both DR and DIO dams. The meal patterns of HE-fed dams, who consumed significantly larger meals, echoes what we previously observed in leptin receptor-deficient rats [29]. Even though the F0/F1 dams largely compensated for the increased caloric density of the HE, showing no differences in 24-hour caloric intake except during late pregnancy, a phenotype effect on food intake was detected in the F3 generation; DIO dams on chow consumed slightly more kcal than DR on chow, confirming our selective breeding methods. An overall diet effect and time x diet interaction was found in dams on HE, who consumed more kcal than animals on chow. These findings are interesting in light of varying postpartum weight change patterns between chow- and HE-fed dams. Despite the relatively similar postpartum caloric intake, dams consuming HE maintained a stable body weight during lactation whereas chow-fed dams exhibited dramatic weight loss followed by regain over the period of lactation. Though we did not evaluate it directly in our studies, one reason for this could involve divergent changes in postpartum energy expenditure. In an earlier study by Whalig and colleagues, mice maintained on HFD before and after pregnancy exhibited reduced total body energy expenditure on postpartum day 10, which in combination with increased caloric intake, induced a state of greater positive energy balance in these HFD-fed dams compared to dams fed chow [13]. The authors of this study went on to demonstrate that in dams consuming HFD, dietary fat is used for milk lipid production, rather than energetically costly de novo lipogenesis, thus contributing to lower total energy expenditure [13]. Furthermore, as leptin signaling promotes energy expenditure, it is reasonable to hypothesize that leptin resistance, as observed in the HE-fed dams in our study, also plays a role in reduced postpartum energy expenditure.

We further speculate whether a state of postpartum leptin-resistance contributes to the consistent, and often severe, liver steatosis that was observed in dams fed HE diet. Deficient leptin signaling reportedly drives lipid accumulation in the liver, as has been observed in hypoleptinemic people with lipodystrophy [49], and in people with type 2 diabetes and leptin resistance [50]. High dietary fat and fructose content are listed as risk factors for liver steatosis in people [51], and mice maintained on a 45% HFD for 24 weeks also displayed liver steatosis [34]. However, our observation that several female rats who failed to become pregnant, but were maintained on the HE for the same duration as the those who did, showed no evidence of liver lipidosis, suggests that the HE diet interacts with the altered metabolic state of pregnancy or lactation to exacerbate lipid accumulation. Interestingly, a similar phenomenon has been described in high-yielding dairy cows. At lactation onset, some dairy cows can experience extreme energy deficit, which triggers hyper-mobilization of lipids from fat stores, thus causing fatty acid release into the blood stream and their accumulation in the liver [52; 53]. This extreme state of negative energy balance in cows results from insufficient food intake following parturition, with a concomitant reduction in plasma leptin levels [54]. Consistent with previous work [46; 55], the chow- and HE-fed rat dams in this study appear to increase their postpartum caloric intake soon after parturition and at a similar rate, yet we propose that exaggerated or sustained leptin resistance in the HE-fed dams creates a perceived state of negative energy balance, culminating in a hyper-mobilization of fat stores akin to a dairy cow. Chronic infusions of leptin to early lactating cows effectively reduced lipid levels in the liver by 28% [56], lending support to a hypothesis that restoring leptin signaling in HE-fed rat dams would also reduce liver steatosis.

### Effects of diet and selective breeding on glucose control and amyiin sensitivity

Overweight and obesity in women is also a known risk factor for gestational diabetes mellitus [57]. Therefore, we investigated the insulin levels and oral glucose tolerance (OGTT) in the polygenic obesity model in during pregnancy and lactation in F0/F1 dams. While we unsurprisingly found no robust indication of gestational diabetes mellitus, we could observe slower glucose clearance in HE-fed dams in F0/F1, during both pregnancy and lactation. Fasting insulin and insulin levels in response to glucose were numerically lower in the HE-fed dams in the postpartum period, which could point to mild beta-cell dysfunction, however these differences failed to reach statistical significance. In the generations F6/F7, a main effect of phenotype was observed during the postpartum OGTT, with DIO on HE demonstrating a delayed glucose clearance compared to DIO-chow dams. To date, most rodent models of gestational diabetes mellitus do not fully recapitulate the human condition—many rely on the use of diabetogenic drugs or manipulation of maternal diet [58; 59], and few studies have investigated the impact of gestational diabetes on maternal health beyond the postpartum period [60]. The data presented here suggest that the selectively-bred DIO model of maternal obesity is not a suitable model of gestational diabetes. However, with further modifications to the maternal diet, such as increasing diet duration or fat or carbohydrate amount or composition, it may present a favorable foundation for future models.

Like leptin, which rises steadily throughout pregnancy, previous studies showed that circulating amylin levels also increase during gestation [61]. And while the occurrence of gestational leptin resistance during pregnancy is established [62], it was not known if this rise in amylin levels also leads to reduced amylin sensitivity, thus providing an additional mechanism to promote increased caloric intake during pregnancy. The treatment of pregnant rat dams with amylin followed by food intake measurements demonstrated that amylin sensitivity is preserved during pregnancy. The effectiveness of amylin to reduce food intake was also not influenced by the diet consumed by the pregnant rats. This finding is consistent with our earlier results, which showed in male rats that obesity or HFD-induced hyperamylinemia did not cause a state of amylin resistance [39].

### Effects of diet and selective breeding on behavioral profile of postpartum dams

In our first experiments in F0/F1 dams, our lower resolution, real-time analysis of maternal behaviors during a pup-retrieval test following by 60 minutes of home-cage observation, failed to detect any effect of HE on maternal behaviors. There was enough variability in these data, however, that left us questioning whether these measures were adequately sensitive to detect potentially subtle changes based on diet. One key aim of the F3 experiment was to perform a high-resolution, blinded analysis of a video-monitored maternal behavior test. Under these experimental conditions, and after three rounds of selective-breeding for obesity-proneness or -resistance, we observed a significant delay in latency to begin nursing in DIO dams, and two outliers within the DIO dam group that demonstrated low pup-related behaviors on P2 (See Figs. 4C and D). When the test was repeated on P7, when maternal behaviors are consolidated, there were no differences between groups. While these subtle differences are consistent with other published findings in mice that HFD can reduce the quality of maternal care [14; 15; 17], another study in rats showed that HFD increased the time that the dams nursed their pups [16]. Because the methods and endpoints used to assess maternal care are not consistent across these or our experiments, it is difficult to determine what are true effects of maternal diet or obesity on maternal care, and what is simply the reflection of natural variations in maternal care that occur over and within the early postpartum days. To bypass such hurdles would require multiple tests, if not constant home-cage monitoring, of maternal care. Further, because the home-cage retrieval test is relatively simple for the dams to complete, this again introduces a question of sensitivity. Incorporation of a novel or stress-evoking environment, such as a T-maze retrieval test [63; 64], would further allow us to assess if maternal diet or obesity influences susceptible or resilience in the face of such a stressor. However, putting these points aside, our blinded analysis of maternal care in F3 dams, suggest that while a polygenic predisposition for obesity might increase the tendency for reduced maternal care, neither intake of HE diet or maternal obesity per se reduced the overall quality of maternal care.

In a previous set of experiments, rat dams exhibiting high maternal care exhibited higher expression of oxytocin in the PVH, and a higher number of oxytocin receptors in brain centers critical for maternal behaviors [65; 66]. Levels of oxytocin can also be influenced by metabolic state. While leptin was shown to increase PVH oxytocin gene expression [67], leptin receptor-deficient rats have reduced plasma oxytocin levels [68]. Based on the observed reduction in leptin-sensitivity in the HE-fed dams, we hypothesized that HE-intake would reduce oxytocin expression in the PVH. The quantification of oxytocin-positive neurons in the PVH of DR and DIO rat dams at the time of pup-weaning, however, failed to detect any effect of phenotype or diet on oxytocin levels. During our analysis, we were sensitive to the heterogenous expression pattern oxytocin neurons across the rostro-caudal axis of the PVH [69], only comparing anatomically aligned sections. One important limitation is that we investigated oxytocin immunoreactivity in the PVH, and not the functional release of oxytocin. Interestingly, there is some evidence that leptin can increase activation of oxytocin neurons [70; 71]. We also quantified oxytocin at P26, when lactation demands on the dam are falling, in contrast to previous studies that quantified oxytocin on P6 [65]. So while these data would suggest that metabolic state does not influence late-postpartum levels of oxytocin in PVH, we cannot rule out that metabolic state or leptin sensitivity might influence release of oxytocin into the periphery or at central target sites.

The lactation period has generally been described as anxiolytic [72]. Rodent dams reportedly spend more time in the open arms of an elevated plus maze (EPM), a readout for reduced anxiety, and this adaptation is thought to be a critical component of the new mother’s heightened drive to protect her young. Furthermore, consumption of HFD and weight gain in obesity-prone non-pregnant females was previously shown to increase anxiety-like behavior in EPM [73], leading to the hypothesis that maternal intake of HFD would prevent the lactation-driven reduction in anxiety. As with maternal motivation, there are conflicting data on the effect of HFD on maternal performance in behavioral paradigms testing anxiety-like behaviors. Moazzam and colleagues reported that postpartum mouse dams maintained on HFD spent less time in the closed arms of an EPM than chow-fed dams, which would actually indicate reduced anxiety [15]. Another study by Perani and colleagues showed that HFD prevented lactation-associated anxiolysis, with HFD-fed mouse dams showing an increase in the latency to enter the lit chamber of the light-dark box, and a decrease in the number of lit chamber entries [74]. Similarly, a study investigated the effects of various maternal diets in mice showed that intake of a diet high in fat and branched-chain amino acids reduced time spent in open arms on P8 [14]. In our studies, we failed to see robust effects of diet in the open field test. We did not observe phenotype differences in OFT measures in virgin females naïve to HE, nor any global effects of HE or phenotype when the same females were tested on postpartum day 10.

To round out the behavioral profiling of this maternal obesity model, we also performed a mid-lactation sucrose preference test, as a readout for hedonia or pleasure-seeking, in the dams from the F3 and F6/F7 generations. In generation F3, we did not see statistical differences across groups. However, similar to the outliers observed in the maternal behavior test, we observed a few individual dams in the DIO groups which displayed a possible sucrose aversion (consuming less than 50% sucrose). In generations F6/F7, we observed a main effect of both phenotype and diet, with DR-chow animals showing higher sucrose preference compared to DIO-chow dams. These data are similar to those reported by Moazzam and colleagues, where pregnant mice on chow displayed higher sucrose preference compared to pregnant mice on HFD [15]. Resistance of the DIO rat dams to consume sweet solution was also observed during the OGTTs performed in the F6 generation, by which we had to abandon the more clinically- and physiologically-relevant drinking of glucose during the OGTT for gavage administration. The DIO rats, despite an overnight fast prior to the test, simply would not drink the glucose solution. While a reduction in preference for sucrose is a commonly used correlate for an anhedonic or depressed state, it should not be neglected that the test also depends on sucrose detection and post-ingestive learning mechanisms [75; 76]. Whether selective breeding based on proneness to weight gain also selects for reduced sucrose preference or detection, in both males and females, would be interesting to investigate. Supporting this, an earlier study demonstrated that obesity-prone male rats maintained on chow demonstrate reduced oral sensitivity to sucrose [76].

### Translational value of the DR/DIO model of maternal obesity

Herein, we have described the effects of a palatable HE diet in two strains of rats during pregnancy: one that is highly susceptible to weight gain, and one that is resistant. We observed that the intake of HE diet during pregnancy and lactation can have robust effects on various readouts of metabolic health in both strains, and that a polygenic susceptibility to weight gain has an additional impact on some of these measures. Because obesity in women is known to increase the risk of various disorders during and after pregnancy, the polygenic- and diet-sensitive nature of this rodent models provides a multi-faceted tool to dissect how diet, genetics and obesity contribute to these adverse health outcomes. Nonalcoholic fatty liver disease (NAFLD), for example, is more common in women with a BMI above 30 at the start of their pregnancy [77]. Our data would suggest that maternal diet contributes more toward liver steatosis than maternal weight gain or polygenic sensitivity to the diet. The current model would be useful to delineate which components of the HE diet have the largest impact on the liver, and whether HE intake during just pregnancy or just lactation would also produce differential outcomes. Postpartum depression (PPD) is a devastating and complicated human condition, which also has clear associations with maternal obesity [10; 78]. While we feel it is important to emphasize that this model of maternal obesity is not a rodent model of PPD, it can be used as a neurobiological tool that will allow us to probe how obesity-induced changes in the brain could increase susceptibility for PPD. The behavioral profiling of DIO dams suggests that neither maternal obesity nor consumption of a palatable HE universally cause PPD-like behaviors in the postpartum dam. Still, it is interesting to the note that a subset of DIO and/or HE-fed dams often showed more aberrant behaviors in the maternal tests and SPT, denoting potential “high risk” dams within these groups. Our findings therefore support a more general idea that neither intake of HE diet nor maternal obesity cause PPD, rather it is the polygenic predisposition to obesity, which may be compounded by a palatable, calorically-dense diet, that increases the risk for developing PPD. The present rodent model is particularly well-suited to address whether susceptibility for obesity and aberrant postpartum behaviors involve deficits in common neural pathways, such as those involving motivation or stress-responsiveness. This model would also help elucidate what influence certain obesity-related changes (like elevated leptin or leptin resistance) have on the levels of and receptivity to gonadal, placental and metabolic hormones, which are changing so rapidly in the peripartum period.

### Conclusion

In summary, we conclude that intake of HE diet during and after pregnancy can influence maternal physiology and behavior in rat dams and that selective breeding based on the sensitivity to this diet can have both independent and compounded effects on maternal health. Further our results demonstrate, that HE or propensity to gain weight does not cause major deficits in maternal behavior, but the presence of “outliers” within the DIO or HE-fed groups suggests that these are risk factors for alterations in maternal mood or motivation. Our model of maternal obesity, which incorporates both HE diet and polygenic susceptibility for weight gain, represents a useful tool for further probing the biological links between maternal metabolic, reproductive and mental health.

## Acknowledgements

We gratefully acknowledge all the technical support from our lab mates who contributed to the development and maintenance of the DR and DIO colonies: Samira Sigron, Sydney Pence, Faith Slubowski, Luca Papini, Sarah Benz, and lastly Christelle Le Foll for her extensive knowledge on establishing these colonies. We also thank our lab mates who assisted with sample and data collection and discussions of our findings: Salome Gamakharia, Marissa Schraner, Lavinia Boccia, and Bernd Coester. We acknowledge the support of Paulin Jirkof for all of her efforts to re-home the surplus rats. And we gratefully acknowledge Thomas Lutz for his support of this project and review of the manuscript.

## Author Contribution Statement

Conceptualization: CB; Methodology; AL, JB, JM, CB; Formal Analysis: AL, JB, CB; Investigation: AL, JB, JM, CB; Writing - Original Draft: AL, JB, CB; Writing - Review & Editing: AL, JB, JM, CB.

