## Supplemental Material for "Influence of high energy diet and polygenic predisposition for obesity on postpartum health in rat dams"

**Supplemental Methods and Results**

### Home-cage Maternal Behavior Analysis (F3 Dams)

#### Extended Methods

On P2 and P7, a one-hour maternal behavior test was recorded with a video camera. (RaspPi Camera, day/night infrared webcam). Starting two hours before the dark phase, litter and dams were weighed. Prior to testing, plastic tubes and some of the nesting material were removed in order to better observe the behavior on the video. Cages were moved to the corresponding rack for the video recording and nest position was noted. The video was always taken from the back of the rack, and cages were placed in rack with the nest at the back. Before lights-off (at 11.00 am), the camera was installed, and 3 red light lamps were positioned to illuminate the video.

Fifteen minutes after lights off, pups were removed from their nest and separated from the dam in a separate cage for approximately 10 minutes. Video recording was started. Pups were then returned to the two furthest corners from the dam's nest (5 pups in each corner). After returning all the pups into their cages, the observer left the room. Video recording was stopped one hour after the last pups had been returned to their cages; plastic tubes and nesting material were returned to the corresponding cages.

The videos were analyzed using the Behavioral Observation Research Interactive Software (BORIS), a free and open-source software [1]. The following behaviors were analyzed: latency to retrieve first pup, latency to retrieve all pups, nursing duration, nursing positions (hover, low crouch and high crouch), nest building, pup grooming, self-grooming, eating, drinking, exploring and quiet (out of the nest). For further definition of the behaviors see Supplemental Table 1. To distinguish the difference between hover, low crouch and high crouch, the criteria defined by Stern & Johnson [2] were used. A screenshot of one of the maternal behavior videos (Supplemental Figure 1) provides examples of the three nursing positions.

A bout of an individual behavior was initiated after 2 seconds of sustained behavior, except for nursing, which had to be displayed for a minimum of 5 seconds in order to be counted as a new behavior. Within BORIS, each behavior was assigned as either a point or state event; a point event is a behavior without duration and a state event is defined as a behavior with a defined start and endpoint signifying a duration over time. The behaviors were divided into pup-directed behaviors and self-directed behaviors. Retrieval time, grooming pups, nursing and nursing position accounted for the pup-directed behavior, whereas self-directed behavior included self-grooming, eating, drinking, exploring and quiet. In some cases, observance of certain behaviors excluded the possibility that other behaviors were observed. For example, if the behavior eating was observed, all the other behaviors were excluded. If the animal changed from eating to exploring only one key, in this case x, had to be pressed to simultaneously end eating and start exploring; the endpoint of eating was then automatically added one millisecond before the start of exploring behavior was logged by the BORIS program. Output data was directly plotted with BORIS and analyzed using Microsoft® Excel and Prim 8.

The observer coding the videos was blinded to the identity of the dam. After rating all videos from P2 and all videos from P7, the first two videos from the corresponding groups were reevaluated to control for an observer reliability. For each of the 11 behaviors scored, there was between 0 and 2% difference in duration between re-scored videos.

| Behavior | Definition | Category |
|---|---|---|
| Retrieval time of first pup | Time from adding pups until the retrieval of the first pup to the nest | No category |
| Retrieval time of all pups | Time from adding pups until last pup was retrieved to the nest | Pup-related |
| Total nursing time | Time spent nursing on the nest. Behavior started after rat settled on the nest for more than 5 seconds | Pup-related |
| Hover | Time spent nursing, while other behaviors such as self-grooming, pup grooming and nest building were displayed | Pup-related |
| Low crouch | Time spent nursing while dam was quiet, moving only to change position and not showing a high arch position | Pup-related |
| High crouch | Time spent nursing while dam showed a high arch position and / or pups were moving without dam moving. Usually behavior displayed following low crouch. | Pup-related |
| Nest building | Time dam spent building the nest. Could be during the hover position on the nest or while the dam is freely moving in the cage and digging nesting material towards the nest. | Pup-related |
| Pup-grooming | Time dam spent grooming the pups | Pup-related |
| Self-grooming | Time dam spent grooming herself. Could be on the nest while nursing and hover or anywhere else in the cage. | Self-related |
| Eating | Time dam spent eating in the corner with the food hopper | Self-related |
| Drinking | Time dam spent drinking from the water bottle | Self-related |
| Exploring | Time dam spent exploring, walking in the cage, sniffing. | Self-related |
| Quiet | Time dam spent sitting/sleeping in another place than the nest without displaying any of the above defined behaviors. | Self-related |

**Supplemental Table 1:** Definition of maternal behaviors.

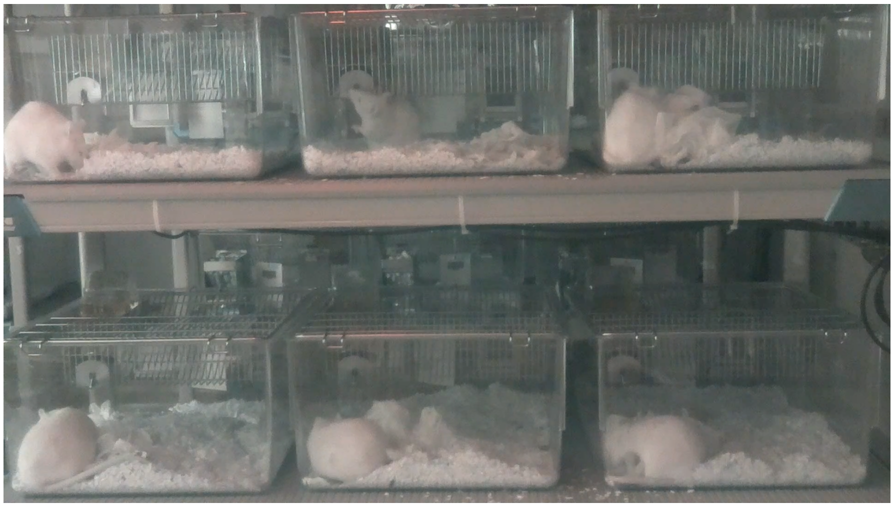

**Supplemental Figure 1:** Screenshot obtained from scored video of maternal behavior test with different nursing positions displayed. The dam in the top left corner exhibits a high nursing position. The dams in the bottom left and bottom middle cages are displaying hover behavior. The dam in the top right corner displays a hovering position, while she is building the nest. And the dam in the bottom right displays a low crouch nursing position.

### Extended Results

On P7, there was a main effect of diet ($F_{1, 24}$ = 5.281, P = 0.03) on pup-grooming behavior, with HE-fed dams showing an increased time dedicated to pup-grooming compared with chow-fed rats, whereas phenotype had no effect ($F_{1, 24}$ = 0.02550, P = 0.87; Supplemental Figure 2D). For all the other behaviors, regardless of postpartum day 2 or 7, there were no further statistical differences between the different groups. Neither phenotype nor diet influenced the maternal behavior on P2 or P7; the complete dataset is shown in Supplemental Figure 2.

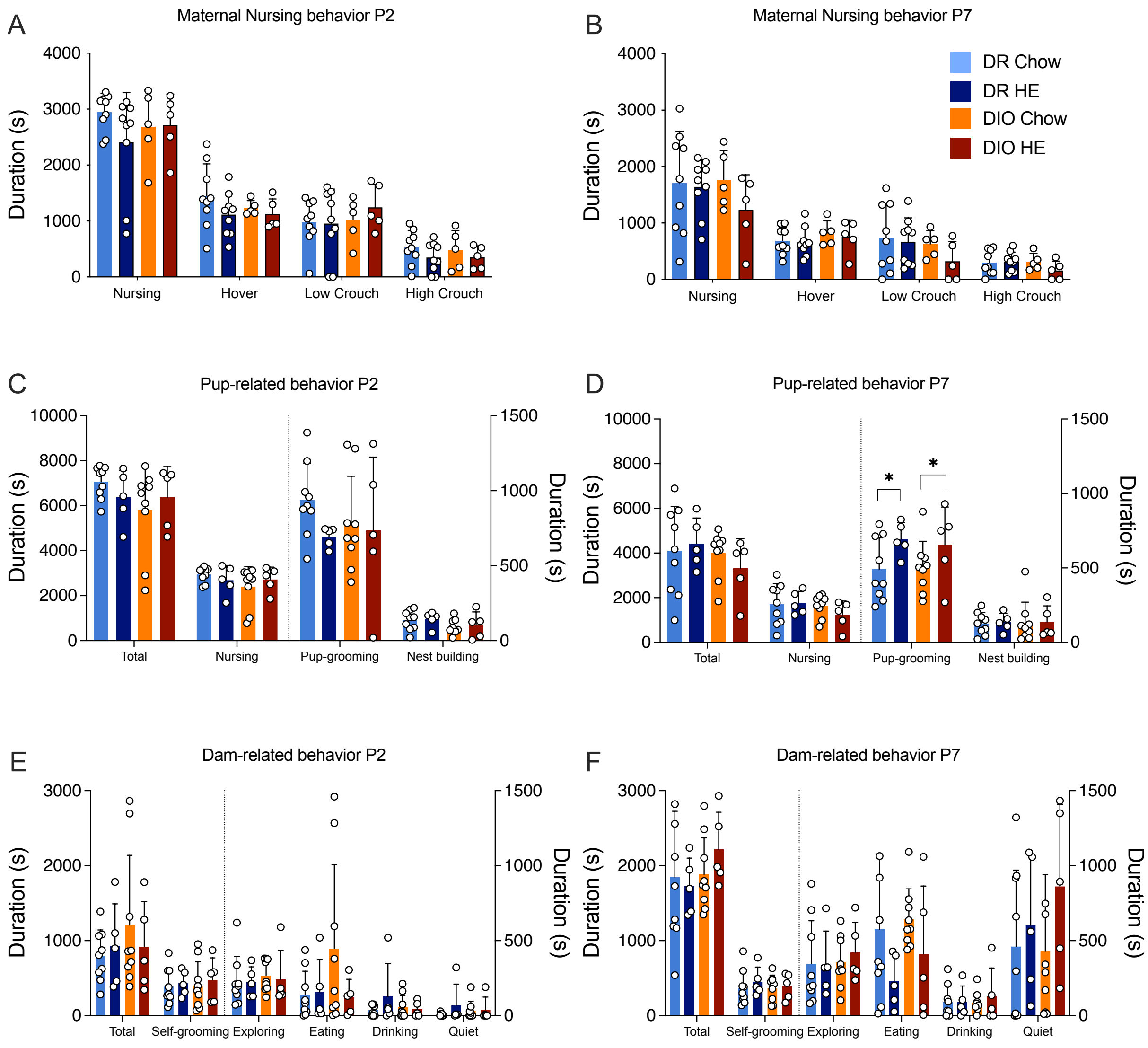


**Supplemental Figure 2: Nursing, pup-related and dam-related behaviors on postpartum day 2 and 7.** (A-B) Pup-related behaviors such as total time nursing, grooming pups and nest building displayed by diet-resistant (DR) and diet-induced obese (DIO) rat dams either on chow or high energy diet (HE). (C-D) Dam-related behaviors including total time self-grooming, exploring, eating, drinking and quiet displayed by DR and DIO rats on either chow or HE diet. (A-D) Data are represented as mean ± SD, symbols denote significant differences between DR vs. DIO chow or HE fed rats: *P ≤ 0.05

**Litter parameters (Offspring of F3 Dams)**

#### Results

To determine if differences existed between litters born to DR or DIO dams, litter size, male to female ratio and litter weight pre-culling were measured. Starting from P2, litter weight was measured daily to assess litter weight gain and individual pup weight gain. Data were analyzed using two-way ANOVA for litter size, male to female ratio and litter weight pre-culling. For litter weight, litter weight gain and individual pup weight gain additionally a mixed effect analysis was used, followed by Tukey's multiple comparisons test.

No significant differences between phenotype ($F_{1, 25} = 1.767$, $P = 0.20$) or diet ($F_{1, 25} = 0.1223$, $P = 0.73$) were found in the litter size (Supp. Figure 3A), nor for male to female ratio (phenotype: $F_{1, 25} = 0.4782$, $P = 0.50$; diet: $F_{1, 25} = 0.005113$, $P = 0.94$; Supp. Figure 3B). Furthermore, there was neither a phenotype ($F_{1, 25} = 2.057$, $P = 0.16$) nor a diet ($F_{1, 25} = 1.014$, $P = 0.32$) effect in the litter weight before culling the litters to ten pups (Supp. Figure 3C).

Regarding the litter weight from day P2 to P25, a time effect ($F_{1.761, 43.57} = 6545$, $P < 0.0001$), a phenotype effect ($F_{1, 25} = 6.448$, $P = 0.0177$) and a time x phenotype interaction ($F_{23, 569} = 3.928$, $P < 0.0001$) was detected. Further, a time x phenotype x diet interaction could be found ($F_{23, 569} = 2.282$, $P = 0.0007$; Supp. Figure 3D). From P7 to P24 there was a significant difference between the DR chow-fed and the DIO chow-fed litter weights (average $P = 0.01$) showing that DIO chow-fed litters are heavier than DR chow-fed litters. At P16 and P17, DR HE-fed litters were significantly lighter than DIO chow-fed litters (average $P = 0.045$).

Analysis of litter weight gain revealed a main effect of time effect ($F_{5.653, 139.6} = 228.2$, $P < 0.0001$). No phenotype or diet effect and no interaction of the factors was observed (Supp. Figure 3E)
In the multiple comparison analysis, DIO chow-fed litters gained more weight compared to DR chow-fed litters on P9 up to P11 and on P14 (average $P = 0.03$). DR-Chow litters were significantly lighter than DIO-HE litters on P13 ($P = 0.02$) and DIO chow-fed litters gained significantly more weight than DIO HE-fed litters on P11 ($P = 0.02$).

To account for the loss of two pups in two different DIO litters, pup weight gain was corrected for the precise number of pups per litter. Following this correction, a time effect ($F_{4.330, 107.5} = 155.1$, $P < 0.0001$) and a phenotype effect ($F_{1, 25} = 8.824$, $P = 0.0065$) was observed with no significant interaction of the parameters (Supp. Figure 3F). DIO chow-fed pups displayed greater weight gain than DR chow-fed pups reached significance on P5, P9 to P11 and P14 (average $P = 0.024$). In addition, DR chow-fed pups gained less individual weight compared to DIO HE-fed pups on P7 and P13 (average $P = 0.016$). On P10 DIO chow-fed pups gained more individual weight compared to DR HE-fed pups ($P = 0.05$).

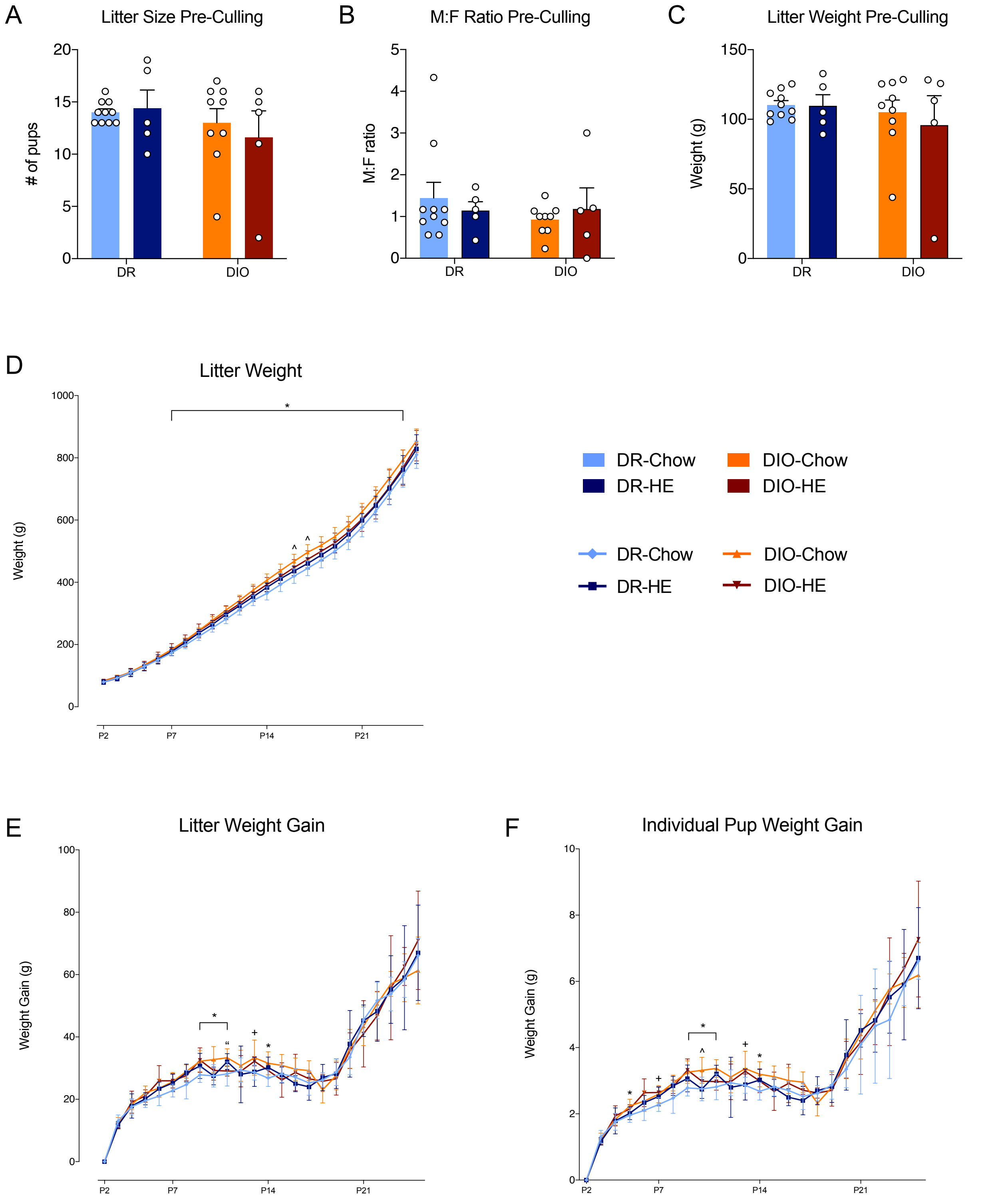


**Supplemental Figure 3: Litter parameters, litter weight and weight gain from postpartum day 2 until 25.** A) Litter size pre-culling on postpartum day 2 (P2) of litters born to diet-resistant (DR) or diet-induced-obese (DIO) rat dams on either chow or high energy (HE) diet. B) Male to female ratio of newborn litters of DR and DIO dams on chow or HE diet on P2 before culling. C) Litter weight of DR and DIO dams on either chow or HE diet before culling on P2. D) Litter weight from P2 to P25 of DR and DIO dams on either chow or HE diet E) Litter weight gain of DR and DIO dams on either chow or HE diet from P2 to P25. F) Individual pup weight gain per litter from DR and DIO dams on either chow or HE diet from P2 to P25. (A-F) Data are represented as mean ± SD, symbols denote significant differences in either litter weight, litter weight gain or individual pup weight gain. * DR-Chow vs. DIO-Chow; + DR-Chow vs. DIO-HE; ^ DR-HE vs. DIO-Chow; " DIO-Chow vs. DIO-HE.

### Open Field Test (F3 Dams)

#### Extended Results

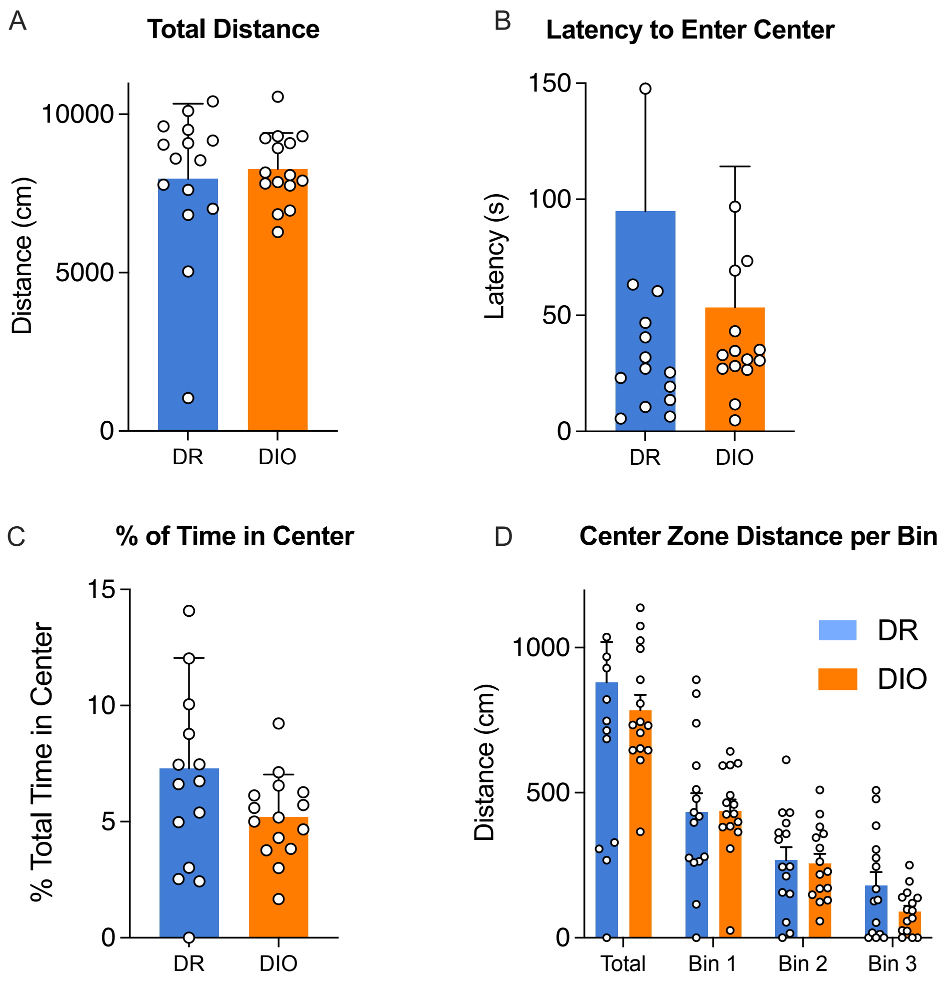


**Supplemental Figure 4: Open field test baseline before gestation in diet-induced obese (DIO) and diet-resistant (DR) female rats on chow diet.** A) Total distance travelled (cm) in session. B) Latency to enter the center zone in seconds. C) Percentage of total time spent in the center zone. D) Distance travelled in center zone per bin. Bin 1 = 0-5 min, Bin 2 = 5-10 min, Bin 3 = 10-15 min. Data are represented as mean ± SD.

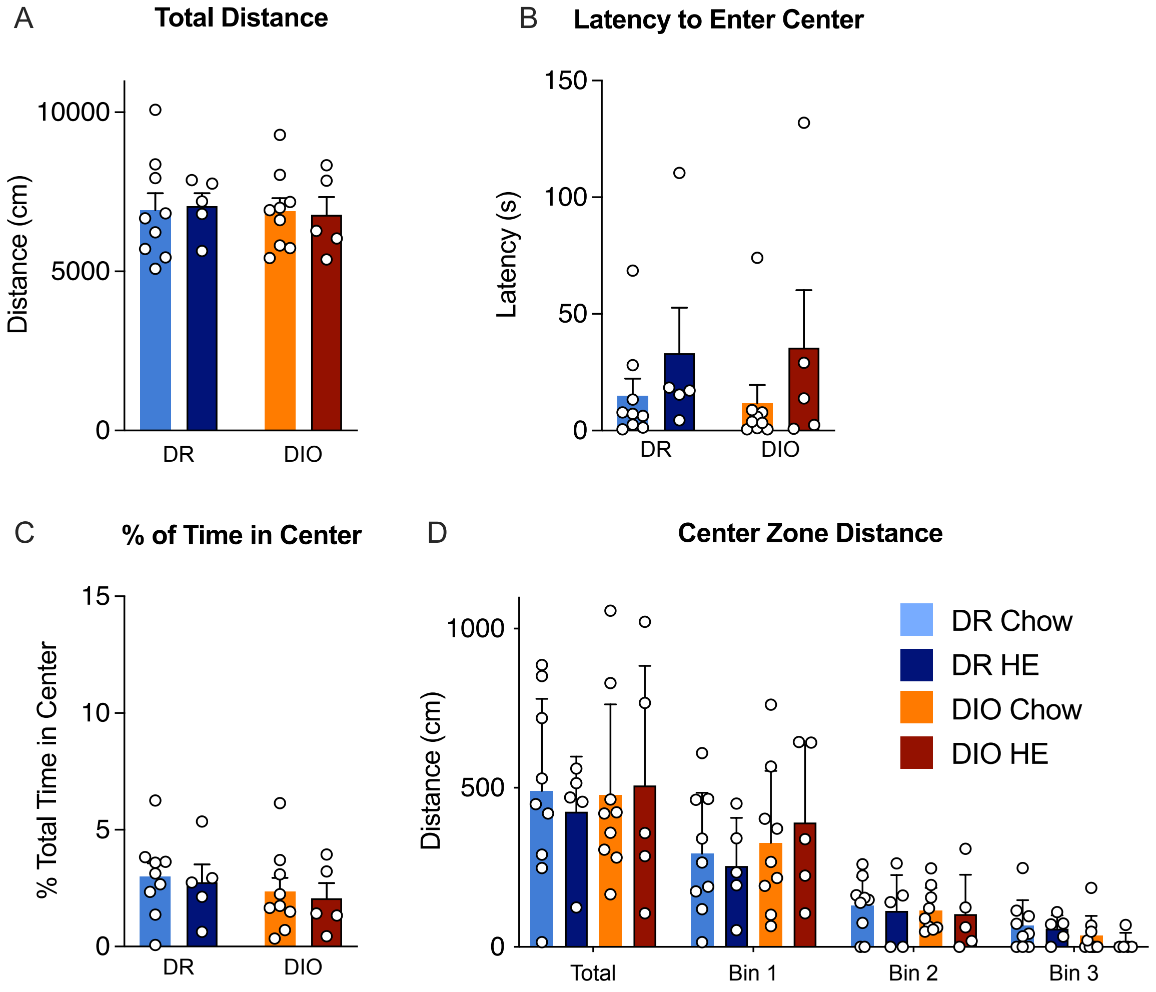


**Supplemental Figure 5: Open field test on postpartum day 10 or 11 conducted with diet-resistant (DR) and diet-induced obese (DIO) rat dams fed either chow or high energy diet (HE) diet.** A) Total distance travelled (cm) in session. B) Latency to enter the center zone in seconds. C) Percentage of total time spent in the center zone. D) Distance travelled in center zone per bin. Bin 1 = 0-5 min, Bin 2 = 5-10 min, Bin 3 = 10-15 min. Data are represented as mean ± SD.

### Quantifying lipid content in postpartum liver

#### Methods

Following semi-quantitative scoring, lipid content of the liver sections from F3 and F7 dams was quantified using the image analysis software Visiopharm (version 2020.08.1.8403; Hoersholm, Denmark). In a first application a Decision forest method was used to outline the whole tissue section as region of interest (ROI). Vessels with a total area greater or equal 17'000 square micrometer were removed from the ROI manually. Finally, a second application with a threshold segmentation method allowed the classification of lipid vacuoles (represented as colorless spaces). Results were represented as total area of lipid vacuoles relative to the total area of liver section.

#### Results

After quantifying lipid content of HE stained liver slides, a diet effect ($F_{1, 23} = 22.46$, $P < 0.0001$) and diet x phenotype interaction ($F_{1, 23} = 6.445$, $P = 0.0184$) was detected (Supp. Fig. 6A). Tukey's multiple comparisons revealed significant differences between DR chow vs. DR HE ($p < 0.001$), DR HE vs. DIO Chow ($p < 0.001$) and DR HE vs. DIO HE ($p = 0.041$), supporting the liver score findings. Further, quantification of the lipid in the F7 livers followed the same pattern, with a main effect of diet ($F_{1,17} = 5.624$, $P = 0.03$; Supp. Fig. 6B), and individual group differences observed between DR-HE and DIO-chow ($P = 0.044$).

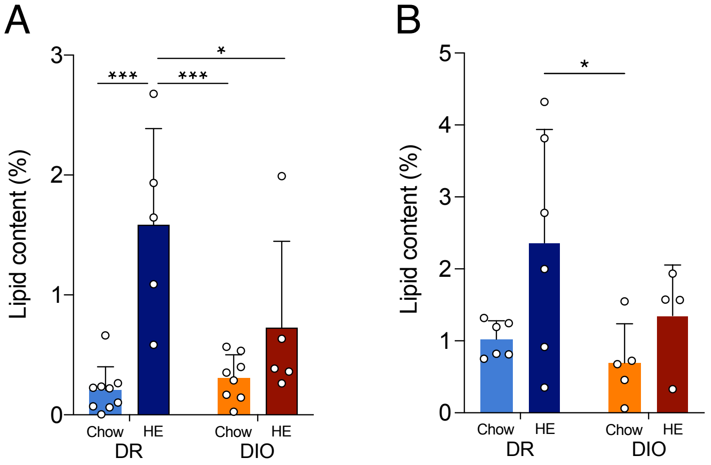


**Supplemental Figure 6: Quantitative evaluation of liver lipidosis using the image analysis software Visiopharm.** Percentage of lipid content in hematoxylin and eosin-stained liver sections in diet resistant and diet-induced obese dams on chow or high energy diet in filial generation F3 (A) and F7 (B).

**Discussion**

Overall, the lipid content quantification confirmed the results from the semi-quantitative scoring, in that dams maintained on HE diet had increased liver lipid content compared to chow-fed dams. The analysis of the F3 livers also revealed a diet x phenotype interaction, reflecting the augmented lipidosis in DR-HE dams. In general, however, the quantitative analysis was limited. Due to widened liver sinusoids caused by increased pressure during the terminal perfusion, the program erroneously identified the sinusoids as lipid droplets, notably in the chow-fed dams which displayed no lipidosis. To utilize this unbiased lipid content quantification method in future experiments, tissue samples will be taken before perfusion in order to avoid dilated sinusoids.
